# Modernizing the NEURON Simulator for Sustainability, Portability, and Performance

**DOI:** 10.1101/2022.03.03.482816

**Authors:** Omar Awile, Pramod Kumbhar, Nicolas Cornu, Salvador Dura-Bernal, James Gonzalo King, Olli Lupton, Ioannis Magkanaris, Robert A. McDougal, Adam J.H. Newton, Fernando Pereira, Alexandru Săvulescu, Nicholas T. Carnevale, William W. Lytton, Michael L. Hines, Felix Schürmann

## Abstract

The need for reproducible, credible, multiscale biological modeling has led to the development of standardized simulation platforms, such as the widely-used NEURON environment for computational neuroscience. Developing and maintaining NEURON over several decades has required attention to the competing needs of backwards compatibility, evolving computer architectures, the addition of new scales and physical processes, accessibility to new users, and efficiency and flexibility for specialists. In order to meet these challenges, we have now substantially modernized NEURON, providing continuous integration, an improved build system and release workflow, and better documentation. With the help of a new source-to-source compiler of the NMODL domain-specific language we have enhanced NEURON’s ability to run efficiently, via the CoreNEURON simulation engine, on a variety of hardware platforms, including GPUs. Through the implementation of an optimized in-memory transfer mechanism this performance optimized backend is made easily accessible to users, providing training and model-development paths from laptop to workstation to supercomputer and cloud platform. Similarly, we have been able to accelerate NEURON’s reaction-diffusion simulation performance through the use of just-in-time compilation. We show that these efforts have led to a growing developer base, a simpler and more robust software distribution, a wider range of supported computer architectures, a better integration of NEURON with other scientific workflows, and substantially improved performance for the simulation of biophysical and biochemical models.

## 1 Introduction

NEURON is an open-source simulation environment that is particularly well suited for models of individual neurons and networks of neurons in which biophysical and anatomical complexity have important functional roles (Hines and Carnevale, 1997). Its development started in the laboratory of John Moore at Duke University in the mid-1980s as a tool for studying spike initiation and propagation in squid axons. Subsequently it underwent massive enhancements in features and performance and it is now used for models that range in scale from subcellular (McDougal et al., 2013) to large networks (Migliore et al., 2006). Today it is one of the most widely used simulation environments for biologically detailed neurosimulations (Tikidji-Hamburyan et al., 2017).

Einevoll et al. (2019) have argued that the central role of simulation software in neuroscience is analogous to physical infrastructure in other scientific domains, such as astronomical observatories and particle accelerators, and that the resources required to build and maintain software should be considered in this context. The increasing importance of software in science is, however, not specific to neuroscience. Crouch et al. (2013) and Hettrick et al. (2014) found that there is a general trend that science relies more and more on software with the capability to automate complex processes and perform quantitative calculations for prediction and analysis. Unfortunately, this reliance on software also has inherent and increasing risks (Miller, 2006). The need for better software sustainability, correctness and reproducibility (McDougal et al., 2016; Mulugeta et al., 2018) has prompted initiatives and proposals suggesting better practices when developing scientific software (Crouch et al., 2013; Erdemir et al., 2020) and when publishing computational results (Heroux, 2015; Willenbring, 2015). In practice, however, it remains difficult to always have the right training, resources and overall understanding to develop good software and use it correctly. Bartlett et al. (2012) and Gewaltig and Cannon (2014) further illustrate how productive use of a software application can lead to development and use beyond its original scope. This, in turn, increases its complexity and can render a once-straightforward implementation unwieldy and hard to maintain.

Another challenge is that of software portability. A user may, rightfully, expect that a scientific software runs on different operating systems and makes good use of all the installed hardware, which in today’s systems often means a combination of a multi-core CPU and a powerful graphics processing unit (GPU). The number and diversity of these hardware architectures is expected to continue to increase as hardware architects seek to further exploit problem specificities in their designs (Hennessy and Patterson, 2017). The increasing difficulty in miniaturizing transistors will amplify the trend towards architectural heterogeneity (Hennessy and Patterson, 2019). From a software point of view, maintaining portability for this diversity of platforms is a fundamental challenge. The more target platforms that need to be supported, the bigger the risk that this leads to multiple redundant code segments with potentially different programming syntax, compilation configurations, and deployment mechanisms, which are error-prone and labor-intensive to maintain. In the software development world, mechanisms and paradigms have been found that facilitate writing more portable software, such as programming paradigms that support multiple architectures (Wolfe, 2021), and modern continuous integration mechanisms (Meyer, 2014). If we want to be able to keep benefiting from future hardware developments in neuroscience, neurosimulator software will have to fully engage with the portability challenge.

Another important challenge is running computational models quickly and efficiently, including those of large size. This requires understanding the computational nature of the scientific problem, which computer system is best suited (Cremonesi et al., 2020; Cremonesi and Schürmann, 2020), and optimization of data structures and algorithms for specific hardware architectures (Kumbhar et al., 2019; Jordan et al., 2018). Given the multitude of computational models and diversity of computer architectures, it has become necessary to use various automated approaches to generate optimized versions of the software. Examples include modern compiler techniques for code generation from domain specific languages (e.g. Blundell et al. (2018); Akar et al. (2019); Kumbhar et al. (2020)), or just in time compilation (Lam et al., 2015), as well as the use of platform-optimized libraries (e.g. Agullo et al. (2009); Carter Edwards et al. (2014)) and new abstraction layers that anticipate heterogeneous architectures (Beckingsale et al., 2019).

A major confounding factor to the aforementioned challenges is that popular scientific codes, such as NEURON (Hines and Carnevale, 1997) or NEST (Gewaltig and Diesmann, 2007), have often been developed over long periods and include key source code that was written without the benefit of modern development tools, libraries and software programming practices. The necessity of modernizing scientific codes is increasingly recognized (de Verdière, 2020; Neely et al., 2017) and does not spare brain simulator software projects (Brette et al., 2007). In the case of NEST these modernizations happened over the past few years, spanning a wide range of both algorithmic and technical improvements (Pronold et al., 2022). Others, such as the Brian Simulator (Goodman, 2009), have decided to rewrite their codes from scratch, taking the opportunity to overcome limitations of their previous implementations, such as allowing for flexibility in model specification while improving simulator performance (Stimberg et al., 2019). It also prompted the inception of new simulator projects, such as the Arbor simulator (Akar et al., 2019), where the developers sought to start from a design philosophy that prefers standard library data structures, code generation, and which minimizes external dependencies. The flip side of such a fresh start is that it is difficult to maintain full backward compatibility with existing models.

Lastly, with more complex scientific workflows, the notion of software being a single do-it-all tool is slowly waning and one should rather think of it as a building block in a larger eco-system. This trend can be seen in efforts like the EBRAINS research infrastructure^1^, where multiple tools are combined into intricate scientific workflows (Schirner et al., 2022). The Open Source Brain (Gleeson et al., 2019) and tools such as NetPyNE (Dura-Bernal et al., 2019), LFPy (Lindén et al., 2014), Bionet (Gratiy et al., 2018) and BluePyOpt (Van Geit et al., 2016) use NEURON as a library and augment it with additional features. As an example, NetPyNE is a high-level Python interface to NEURON that facilitates the development, parallel simulation, optimization and analysis of multiscale neural circuit models.

Here, we report on our efforts to modernize the widely-used NEURON simulator, improving its sustainability, portability, and performance. We have overhauled NEURON’s overall code organization, testing, documentation and build system with the aim of increasing the code’s sustainability. We have integrated NEURON with an efficient and scalable simulation engine (CoreNEURON) and a modern source-to-source compiler (NMODL) capable of targeting both CPUs and GPUs. We demonstrate the performance of several large-scale models using different multi-CPU and multi-GPU configurations, including running simulations on Google Cloud using NetPyNE. Finally, we updated NEURON’s reaction-diffusion simulation capabilities with just-in-time compilation, more seamless integration with the rest of NEURON, support for exporting to the SBML format, and support for 3D intra- and extra-cellular simulation.

## 2 Methods

Over the years the NEURON simulator has been developed to accommodate new simulation use-cases, support community tools and file formats, and adopt new programming paradigms to benefit from emerging computing technologies. During this process, various software components have been developed and external libraries have been integrated into the codebase. Like many scientific software packages, maintaining a codebase developed over four decades poses a significant software engineering challenge.

Since the 7.8 release the NEURON developer community has launched a variety of initiatives to future-proof the simulator codebase. Figure 1 summarizes the high-level functional components of the NEURON simulator and the various changes described in the rest of this section. These developments happened since 2020 over the course of two years starting with the refactoring of the build system and the introduction of a new continuous integration (CI) system. While especially the latter is an ongoing effort, both developments have allowed various improvements to documentation, testing and packaging. The tighter integration of CoreNEURON into NEURON and its various performance and hardware portability improvements were implemented in parallel and were able to quickly benefit from the new build system and CI made available in NEURON.

**Figure 1:**
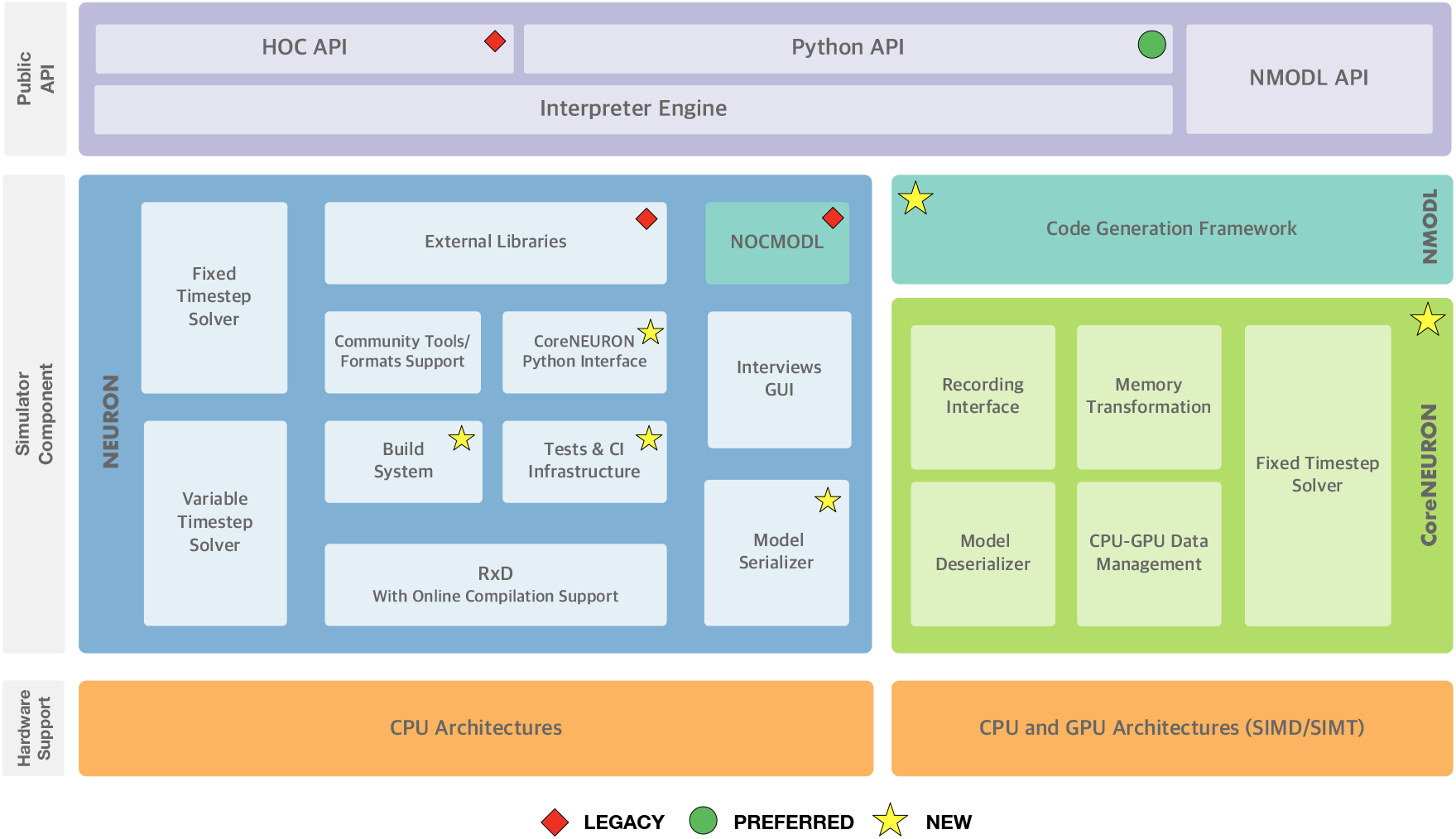
NEURON Simulator Overview: At the top, the “Public API” layer shows NEURON’s three application programming interfaces exposed to end-users: the legacy HOC scripting interface, the preferred Python interface, and the NMODL DSL for defining channel and synapse models. In the middle, the “Simulator Component” layer shows the main three different software components and their interal sub-components: NEURON, the main modelling and simulation environment, CoreNEURON, a compute engine for NEURON targetted at modern hardware architectures including GPUs, and NMODL, a modern compiler framework for the NMODL DSL. At the bottom, supported hardware architectures are shown. Software components that are newly added or are deprecated are highlighted.

### 2.1 Improving software sustainability through code modernization and quality assurance

To address some of the shortcomings of NEURON’s codebase accumulated over four decades of continued development, we have implemented a number of far-reaching changes and adopted a new modern development process with the aim of streamlining the handling of code contributions, while at the same time improving code quality and documentation. First, we have replaced the legacy GNU Autotools based build system with CMake. Second, this allowed us to introduce a comprehensive automated build and CI system using GitHub Actions. Third, these two changes allowed us to create a modern binary release system based on Python wheels. Finally, we have further extended these components to integrate a code coverage monitoring service and automatically build user and developer documentation.

#### 2.1.1 Modern Build System adoption

Until recently the build system of NEURON used GNU Autotools. Autotools, the de-facto standard on Unix-like systems, is a build system used to assist the various build steps of software packages. CMake is a modern alternative to Autotools that offers many advantages and features important for the continued development of NEURON. First, it has extensive support for customizing C and C++ builds, from language standards, to fine-tuning compile and link-time arguments. Furthermore, it supports build portability across hardware platforms (i.e. x86_64, ARM, GPUs), operating systems (i.e. Linux, macOS, Windows) and compilers (GCC, Clang, Intel, NVIDIA, etc.). It also allows a more robust integration with external dependencies. Finally, CMake is being actively developed and supported by a large community of open source and industry developers.

We decided, therefore, to replace NEURON’s legacy Autotools build system by CMake and reimplemented the entire configure and build process using CMake for NEURON as well as its libraries such as CoreNEURON and Interviews. The build-system reimplementation allowed us to refactor large parts of the auxiliary code used for configuring, packaging and installing NEURON in order to make it more robust and maintainable.

Using CMake we are able to provide a build configuration that goes far beyond GNU Autotools in several respects. For instance, we have included the ability to automatically clone and integrate other CMake based libraries like CoreNEURON and NMODL using CMake options. The build is organized in various build targets producing multiple shared libraries for the interpreter, solvers, simulator, the native interfaces of the Python API, RxD, CoreNEURON, and the main neuron executable, `nrniv`.

#### 2.1.2 Continuous integration and build automation

Continuous integration (CI) is crucial for the development process of any software project. It allows the development team to check the correctness of code changes over the course of the project’s development life cycle. To support our work in modernizing the NEURON codebase and opening up the development process to a wider community it was, therefore, important to first put a CI in place. Figure 2-A gives an overview of the CI workflow using GitHub Actions and Azure. Every time a new *Pull Request* is opened on the NEURON repository, CI pipelines for building and testing NEURON are executed on Linux, Windows and macOS. These also rebuild the documentation, executing all embedded code snippets and generate test code coverage reports, which are provided to the code reviewer to aid them in evaluating the proposed change to the code. As part of the CI workflows, we also need to build and test binary installers as well as python wheels for various platforms. As Github Actions provides limited concurrent builds for open source projects, we use Azure CI workflows for building artifacts such as installers and python wheels. This helps us to reduce overall CI turnaround time.

**Figure 2:**
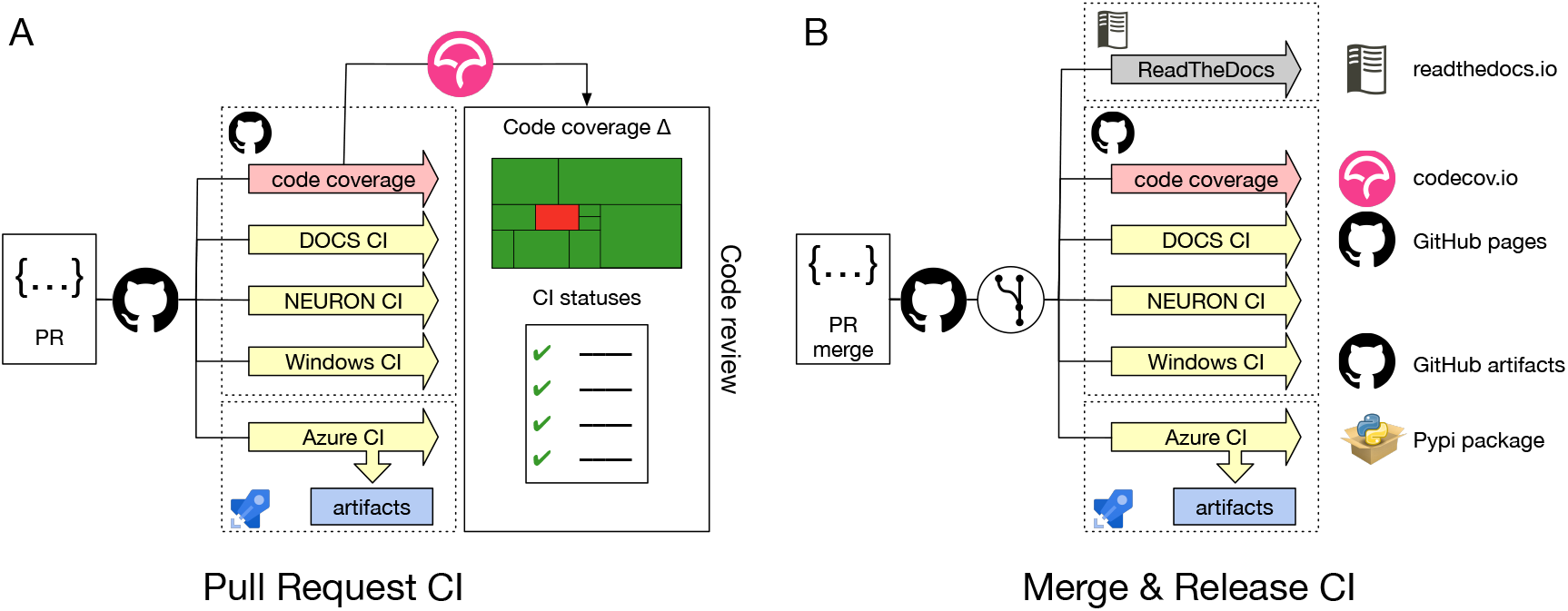
**(A)** *Pull Request* CI workflow: Whenever a *Pull Request* is opened jobs are started to build and test NEURON on Linux, Windows and macOS. Several combinations of build options and versions of dependencies are tested. Also, Python wheels are built, the documentation is regenerated, executing embedded code snippets and code coverage metrics are determined. All CI job results are reported to the code reviewer. **(B)** Merge & Release CI workflow: Automated CI jobs are also started after a PR has been merged, nightly or on a new version release. These jobs produce artifacts that are delivered into appropriate channels

Using the *Pull Request* CI workflow, we ensure that the proposed change will not introduce bugs or unintended sideeffect, by testing the majority of build options along with different compiler versions. The jobs also build Python wheel packages (see Section 2.1.4 for more details) and test the integrity of the resulting packages by running NEURON’s unit and integration test suites. These jobs are executed both on Linux and macOS systems. Because Windows is a substantially different environment, we needed to develop a separate CI to build and test the NEURON installer package using the MinGW compiler and MSYS2 environment^2^.

Contrary to the Pull *Request* CI, the *Merge & Release* CI (Figure 2-B) is executed on the latest master branch of the NEURON codebase or with a new release. Artifacts such as the latest documentation, Python wheels, binary packages and the Windows installer are built.

Once we had set up a robust build system and CI workflow, we were able to extend the framework to offer additional features both for the developer and the user community. First, as the name suggests, the *Docs CI* builds a NEURON Python wheel package and rebuilds the NEURON documentation (described in more detail in Section 2.1.3). Second, we have added a *code coverage* workflow, which builds NEURON with all features enabled and runs all tests tracking code coverage using `lcov` and `pytest-cov`.

#### 2.1.3 Documentation generation

NEURON’s documentation consists of a number of resources covering the Interviews graphical interface, the HOC and Python APIs, the NMODL domain specific language (DSL), and development best practices.

As part of our effort to streamline the development process and modernize the organization of NEURON, we consolidated NEURON’s documentation sources into the main repository and automated the documentation building. Additional resources, such as tutorials, in the form of Jupyter notebooks, were also gathered and integrated into the NEURON documentation.

We integrated documentation building into the CMake build system and CI pipelines. In the CI pipelines, we start by executing the Jupyter notebooks using a freshly built NEURON wheel, ensuring that existing notebooks are compatible with the latest code. Once the notebooks have been successfully executed, they are then converted to HTML. Next, Doxygen code documentation is generated. Finally, the manual and developers guide are re-built with Sphinx, embedding the previously generated Jupyter notebooks and Doxygen. This documentation is then published on the ReadTheDocs^3^ where we provide versioned documentation starting with the NEURON 8.0.0 release.

#### 2.1.4 A modern NEURON Python package

NEURON was originally packaged as a traditional software application, made available as a binary package for mainstream operating systems and alternatively as a source tarball. Alternatively, NEURON could also be installed as a Python package through a laborious multistep process that lacked flexibility and was error prone. With the introduction of a standard, complete Python interface (Hines et al., 2009), NEURON could be more readily used from a Python shell. Thanks to the user friendliness and strong scientific ecosystem of the Python language, this API quickly became popular in the NEURON community. At the same time, the Python wheel package format has become an extremely popular means of distributing Python software packages, allowing the user to install Python packages using `pip install`.

To provide the flexibility of `pip`-based installation, using the CMake build system we implemented a new NEURON Python package shown in Figure 3. This package is comprised of C/C++ extensions providing the legacy `hoc` language, the Python interface, reaction-diffusion module (`rxd`) with Cython extension, and several pure Python modules. To build functional Python extensions it is important to provide the build system with the correct build flags and paths compatible with the target Python framework. To achieve this we extended the `Extension` class of `setuptools`.

**Figure 3:**
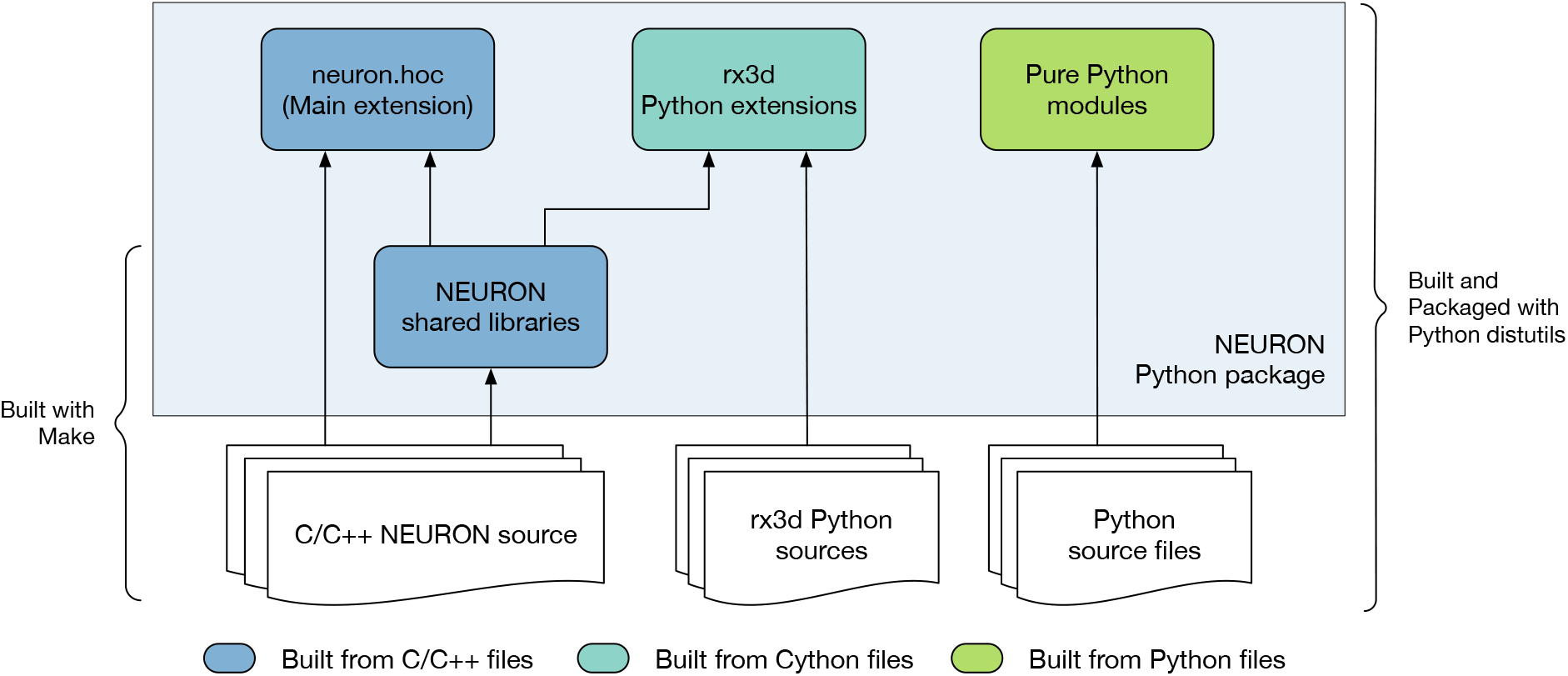
Overview of the NEURON Python package. The package is comprised of pure Python modules and extension code. The main NEURON extension is written in C/C++ and provides the API for the NEURON interpreter. Additionally, the rx3d modules are written in Cython providing the reaction-diffusion solvers of NEURON.

One of the challenges faced when building the NEURON Python package was distribution of binary executables and support for compiling `MOD` files on the user’s machine. Python extensions are typically built as shared libraries and automatically placed in the correct location in the package path to be found at runtime. However, compiled executables such as `nrniv` are treated by `setuptools` as binary data and not installed into the Python framework’s `bin` folder. Also, we needed to support the `nrnivmodl` workflow where the user can compile mechanism and synapse models from MOD files and load the corresponding library at runtime. In order to support this, we created Python shims that take the place of the actual NEURON executables in the `bin` folder. These shims prepare the runtime environment and call the homonymous NEURON binary executable via the `execv` routine, which substitutes the shim process with the NEURON executable.

### 2.2 Integration of CoreNEURON within NEURON

CoreNEURON (Kumbhar et al., 2019) is a simulation engine for NEURON optimized for modern hardware architectures including CPUs and GPUs. CoreNEURON is developed and maintained in its own public repository on GitHub. Previously it was up to the user to obtain API-compatible versions and build NEURON and CoreNEURON from source separately. The use of CoreNEURON was also not straightforward as it had to be run as a separate executable. To simplify the usage, CoreNEURON and NEURON are now coupled more tightly in terms of code organization, build system and implementation level.

First of all, we have integrated CoreNEURON (along with NMODL) as git submodules of the `nrn` repository, allowing us to build single software distribution packages containing optimized CPU and GPU support via CoreNEURON. This also simplifies tracking changes in the various repositories and making sure that the correct code revisions are distributed and built together. Secondly, we have integrated CoreNEURON and NMODL building into NEURON’s CMake build system. This is possible thanks to CMake’s robust support of subprojects that allow easy popagation of build parameters across the various code bases. Thirdly we have implemented an in-memory model transfer, improved GPU support in CoreNEURON, and integrated our `MOD` file code generation pipelines. Some of the important changes are discussed in the remainder of this section.

#### 2.2.1 Transparent execution via CoreNEURON using in-memory model transfer

While CoreNEURON can be run as a standalone application, it still requires the model created by NEURON as an input. This model can be written to disk and transferred to CoreNEURON via files. This approach has the advantage that a large model can be constructed once by NEURON and stored to files, which can be later run by CoreNEURON many times (for example for ensemble runs). Also, this allows the models with huge memory requirement to be constructed in smaller pieces by NEURON before being executed simultaneously by CoreNEURON, thanks to it having a 5 - 6 × smaller memory footprint than NEURON. However, we found that in practice this workflow is not flexible enough for many users.

To address this we have now implemented two-way in-memory data transfer between NEURON and CoreNEURON, which greatly simplifies CoreNEURON usage. With this, it is now possible to record cell or mechanism properties (e.g. voltage, current, variables of type STATE, PARAMETER, RANGE, ASSIGNED defined in MOD files), unlike with the file-transfer mode where only spikes can be recorded. In case of NEURON, all of the data structures representing a model are laid out in an Array-of-Structures (AoS) memory layout. This allows easy manipulation of sections, channels and cells at runtime, but it is not optimal for memory access and Single Instruction Multiple Data (SIMD) execution on modern CPU/GPU architectures. Hence, NEURON data structures are serialized and transferred to CoreNEURON where they are transposed into a Structure-of-Array (SoA) memory layout. This allows efficient code vectorization and favors coalesced memory access, which is important for runtime performance. In addition to data structures representing cells and network connectivity, event queues are now also copied back and forth between NEURON and CoreNEURON, allowing simulations to be run partly with NEURON and partly with CoreNEURON if desired. For end users, all this functionality is now exposed via a new Python module named `coreneuron`. The API and new options are discussed in Section 3.3.

#### 2.2.2 Enabling GPU offloading in NEURON simulations

It is now possible to offload NEURON simulations transparently to GPUs using CoreNEURON. This support is implemented using the OpenACC programming model. When CoreNEURON GPU support is enabled, all data structures representing the model are copied to GPU memory when initializing CoreNEURON. State variables for the Random123 (Salmon et al., 2011) library are also allocated on the GPU using CUDA unified memory. Once the data are transferred, they reside in the GPU memory throughout the simulation. This simplifies memory management and reduces expensive CPU-GPU memory transfers.

All computationally intensive kernels of the main simulation loop are offloaded to the GPU, including state and current updates in `MOD` files, and the Hines solver (Hines, 1984). Even though the spike detection kernels are not computationally expensive, they are offloaded to GPU too, to benefit from data locality and avoid additional data transfers. Only the spike events generated on the GPU and user-requested state variables are copied back to CPU. Similarly, the spikes communicated by other processes are first queued on the CPU and then transferred to GPU.

For some models the Hines solver can consume a significant fraction of the total simulation time when running on GPUs. This is often due to the limited parallelism and non-coalesced memory access arising from heterogeneous, branched tree structures in neuron morphologies. As presented by Kumbhar et al. (2019), two node ordering schemes (called cell permutations) were developed to improve the parallelism and coalesced memory accesses. We have further improved this implementation, reducing solver execution time on GPU by an additional 15 - 20 %. Finally, simulations running on GPU can now utilize multiple GPUs available on a compute node. The available GPUs are uniformly distributed across MPI processes and threads.

#### 2.2.3 Integration of code generation pipelines

An integral part of NEURON’s modeling capability is provided via the NMODL DSL. This allows the user to describe in their models a wide range of membrane and intracellular submodels such as voltage and ligand gated channels, ionic accumulation and diffusion, and synapse models. Such models are written in MOD files and then translated to lower level C or C++ code using a source-to-source compiler (transpiler).

In order to support execution via both NEURON or CoreNEURON, each MOD file is now translated twice: first into C code for NEURON, and then into C++ for CoreNEURON. This workflow is illustrated in Figure 4. `nrnivmodl` remains our main tool for processing user provided `MOD` files. By default, all input `MOD` files are translated into C code by NEURON’s legacy NOCMODL transpiler. These files are then compiled to create a library for NEURON called `libnrnmech`. If a user provides the `-coreneuron` CLI option then either the default MOD2C or the new NMODL transpiler (Kumbhar et al., 2020) is used to translate `MOD` files into C++ files. These files are then compiled to create a library for CoreNEURON called `libcorenrnmech`. The NMODL transpiler generates modern, optimized C++ code that can be compiled efficiently on CPUs or GPUs. These two libraries are finally linked into an executable called `special`. When running on the CPU the user has the choice between using `python, nrniv` or the `special` executable to launch simulations. When running on a GPU, however, one must use the `special` executable to launch simulations due to limitations of the NVIDIA compiler toolchain when using OpenACC together with shared libraries.

**Figure 4:**
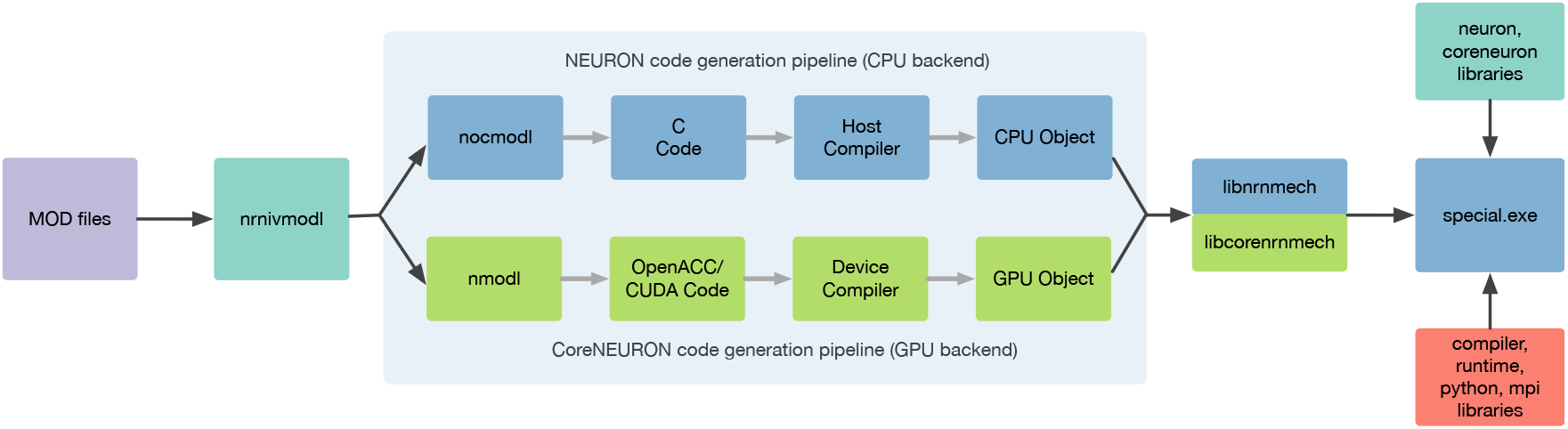
`nrnivmodl` workflow where input `MOD` files are translated into C/C++ code for NEURON and CoreNEURON targeting different hardware platforms via NOCMODL and NMODL. The output of this workflow is a `special` executable containing NEURON and CoreNEURON specific libraries.

### 2.3 Modular NEURON - the example of NetPyNE

NetPyNE (Dura-Bernal et al., 2019) is a high-level declarative NEURON wrapper used to develop a wide range of neural circuit models (Bryson et al., 2021; Pimentel et al., 2021; Volk et al., 2021; Ranieri et al., 2021; Metzner et al., 2020; Anwar et al., 2021; Sekiguchi et al., 2021; Dura-Bernal et al., 2022a,b; Borges et al., 2022; Romaro et al., 2021)^4^, and also as a resource for teaching neurobiology and computational neuroscience.

The core of NetPyNE consists of a standardized JSON-like declarative language that allows the user to specify all aspects of the model across different scales: cell morphology and biophysics (including molecular reaction-diffusion), connectivity, inputs and stimulation, and simulation parameters. The NetPyNE API can then be used to automatically generate the corresponding NEURON network, run parallel simulations, optimize and explore network parameters through automated batch runs, and visualize and analyze the results using a wide range of built-in functions. NetPyNE can calculate local field potentials (LFPs) recorded at arbitrary locations and has recently been extended to calculate current dipoles and electroencephalogram (EEG) signals using LFPykit (Hagen et al., 2018).

NetPyNE also facilitates model sharing by exporting to and importing from the NeuroML and SONATA standardized formats. All of this functionality is also available via a web-based user interface^5^. Both Jupyter notebooks and graphical interfaces are integrated and available via the Open Source Brain (Gleeson et al., 2019) and the EBRAINS (Amunts et al., 2019) platforms.

Simulating large models in NEURON/NetPyNE is computationally very expensive. Thus, enabling CoreNEURON within NetPyNE is very attractive and may provide large gains. Thanks to the tighter integration of CoreNEURON into NEURON (see Section 2.2), we were able to easily integrate CoreNEURON solver support into NetPyNE. We used the `coreneuron` Python module provided by NEURON and added three new configuration options in the simulation configuration object of NetPyNE: `coreneuron` to enable CoreNEURON execution, `gpu` to enable or disable GPU support, and `random123` in order to enable Random123-based random number generators. Users can now enable these features by simply setting the above options in their NetPyNE simulation configuration file.

### 2.4 Enabling new use-cases with reaction-diffusion integration

The NEURON reaction-diffusion module (RxD, McDougal et al. (2013)) provides a consistent formalism for specifying, simulating, and analyzing models incorporating both chemical signaling (chemophysiology) and electrophysiology. Such models are common in neuroscience as, for example, calcium concentration in the cytosol affects the activity of calcium-gated potassium channels. Before NEURON’s RxD module, these models incorporated chemical effects in any of a variety of ways using custom NMODL code; this variation unfortunately made some such models incompatible with each other and posed challenges when combining the custom code with NEURON’s built-in tools. Due to the use of custom solutions, they also generally combined simulation methodology with model description; NEURON’s RxD module, by contrast, explicitly separates the two, allowing, for example, the same model to be used for both 1D and 3D simulation. Recent enhancements to RxD have focused on improving its domain of applicability and usability through changes to the interface and redesigning the backend for more flexibility and faster simulation.

A number of features have been implemented to expand the ability of RxD to better represent a researcher’s conceptual model. An `Extracellular` region type (Newton et al., 2018) provides support for studying cellular interactions through changes in the extracellular space (e.g. in ischemic stroke or between neurons and astrocytes), simulated using a macroscopic volume averaging approach. Three-dimensional intracellular simulation (McDougal et al., 2022) allows study of microdomains and the sensitivity to precise positioning of synapses. Importantly, each of these extensions was designed to fit within the broader RxD context; reaction and diffusion rules are specified and interpreted in the same way for 1D and 3D simulation and for intra- and extracellular simulation. Current through NMODL-DSL-specified ion channels generates a flux in the corresponding extracellular compartment and ionic Nernst potentials are updated based on the 3D extracellular concentration. Both 3D intra- and extra-cellular dynamics are calculated using a parallelized adaptation of the Douglas-Gunn alternating direction implicit method (Douglas and Gunn, 1964). For models not requiring full 3D simulation, it is still sometimes advantageous to account for geometry changes (e.g. in a model using nested shells to account for radial variation, a spine most naturally connects to only the outer-most shell); to allow modelers to address this connection, we added a `MultipleGeometry` to explicitly bridge across geometry changes.

Other interface enhancements focused on extending RxD’s usability. We have worked to make existing NEURON tools work directly with RxD objects. For example, `h.distance` computes path distance between two points, whether they are RxD nodes or segments. Likewise, the NEURON-specific graph types (`h.PlotShape` and `h.RangeVarPlot` for visualizing concentration across an image of the cell and along a path) can plot traditional NEURON variables (e.g. `v` or `cai`) as well as RxD chemical `Species` in whatever region (cytosol, ER, etc.) when using the graph’s matplotlib or pyplot backends. We added a `neuron.units` submodule with conversion factors to facilitate specifying models with rate constants most naturally expressed in specific units (e.g. circadian models involve protein concentrations that change over hours whereas the gating variable on a sodium channel may have a time constant of milliseconds). A new `rxd.v` variable allows using the RxD infrastructure to include dynamics driven by membrane potential, offering an alternative to specifying ion channel kinetics through NMODL files. To allow studying NEURON RxD models with other tools, we introduced `neuron.rxd.export.sbml` which allows exporting reaction dynamics at a point to SBML, a standard for representing systems biology models (Keating et al., 2020).

Additionally, we modified the backend for `h.SaveState` to introduce an extensible architecture for storing and restoring new types of state variables. Python functions within the `neuron` module allow registering SaveState extension types, which consist of specifying a unique identifier, a function to call that serializes the corresponding states, and a function that expands a serialized representation. We implemented an RxD extension that registers itself when RxD states are defined (in particular, importing the module alone does not trigger the registration). When no extensions (including RxD) are present in the model, the saved file is bitwise identical to previous (NEURON 7.x) versions; when extensions are present, the saved data includes a new version identifier, is otherwise identical to the previous version, but ends with binary encoded data representing the number of save extensions used in the model, an identifier for each used save extension, the length, and data to be passed to the extension. In the case of the RxD SaveState extension, as the state data is potentially voluminous, the serialized data is zlib compressed.

We redesigned the RxD backend to improve the flexibility of interactive model specification and debugging. `Region` objects now take an optional `name` argument that can be specified at creation or after to help distinguish them during debugging. For both intra- and extracellular 3D simulation, each `Species, State`, and `Parameter` has its values stored in independent memory locations accessed by a pointer to the corresponding mesh. This architecture allows pointers to be preserved as new `Species`, etc. are defined and old ones are removed. To improve the portability of cells between models, we replaced the requirement that a given species (e.g. calcium) could only be defined once with a requirement that there be no overlapping versions of the same species. This change now allows each cell object in NEURON to fully specify its kinetics, including the reaction-diffusion aspects, thus allowing such cells to be reused in other models without further code changes.

With successive releases of NEURON, we iteratively improved the performance of RxD simulation. We moved all simulation code to C++ and compile reaction specifications to eliminate Python overhead. Reactions continue to be specified in Python as before, but now contain a method that generates a corresponding C file; this method is automatically called when first needed and the corresponding file is compiled and dynamically loaded into NEURON. We replaced the matrix solving algorithm used in 3D RxD simulations with one that exploits the decoupled nature of the reaction-contribution to the Jacobian that arises from reactions only happening between molecules at the same spatial location. Multi-threading support was added to 3D simulations. 3D diffusion rates depend on the concentration at a node and up to six of its neighbors. To minimize cache-miss latency when accessing neighboring voxels when simulating extracellular diffusion,`__builtin_prefetch` was used to move data into a cache before accessing it. For extracellular diffusion, prefetching provides a modest improvement depending of the size of the simulation, e.g. 5% speed-up in a 128^3^ cube of voxels. NEURON does not use prefetching with intracellular simulation, as in practice we observed no comparable speedup. Finally, to accelerate both the initialization and simulation of models with reaction-diffusion dynamics to be studied in full 3D, we now construct voxel based representations of each of its constituent convex components (frusta and their joins) on a common mesh and merge them together (McDougal et al., 2022), instead of constructing a voxel-based representation of an entire neuron morphology at once.

## 3 Results

### 3.1 Sustainability improvements through modern development practices

#### 3.1.1 Towards a development community

As described in Section 2.1, Section 2.1.1 and Section 2.1.2, we have radically updated NEURON’s development life cycle to be a modern and collaborative process. First, new developers are now able to quickly get started thanks to improved documentation (Section 2.1.3). Users can access the latest release documentation at https://nrn.readthedocs.io/en/latest/ and a nightly documentation snapshot at https://neuronsimulator.github.io/nrn/. Second, a modernized build system eases local setup and testing of proposed code changes. A single repository, https://github.com/neuronsimulator/nrn, now provides access to all software components including Interviews, CoreNEURON, NMODL, tutorials, and documentation. The integrated CMake build system across these components provides a uniform interface to build all components with ease. Third, code contributions are automatically checked using a comprehensive CI suite. This increases programmer confidence and helps reviewers to more quickly evaluate proposed changes.

These improvements have directly led to the adoption of a collaborative development process with a lively community. As an example, Figure 5 depicts commits over time since the `nrn` git repository was started in November 2007. We can see that NEURON has been receiving more and more contributions from new developers in the last 3 years.

**Figure 5:**
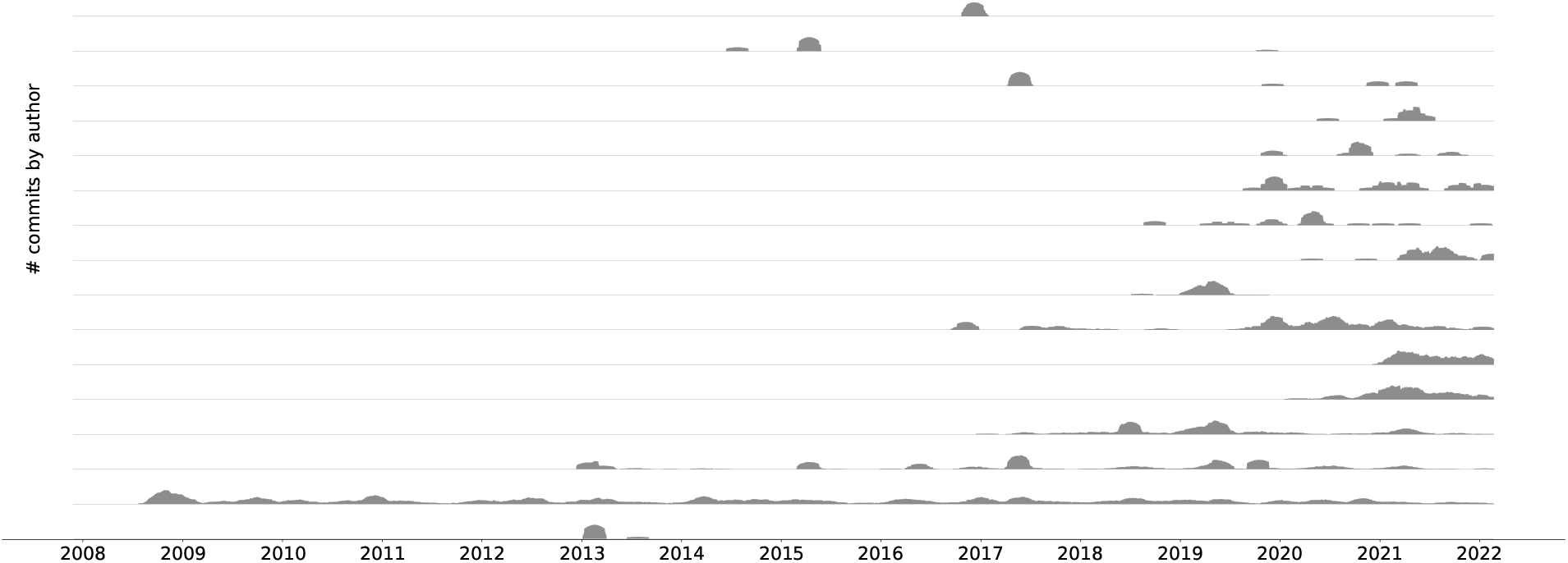
An aligned commit series^6^ plot showing the top 16 contributors since the beginning of the git repository history. For better visibility the different time series do not share y-axes. A clear trend towards collaborative development can be seen.

#### 3.1.2 Software sustainability through development ecosystem modernization

Build system modernization, removing obsolete dependencies, and the introduction of CI pipelines have lead to a vastly streamlined development process. First, using fewer external dependencies sped up and simplified the build process and reduced build times. Second, replacing the Autotools build system with CMake has allowed us to more simply integrate Interviews, CoreNEURON and NMODL in the build and has made build system changes more maintainable. Third, the removal of Autotools and support for legacy Python 2 has simplified the overall code structure, and subsequent code refactoring has become less complex. Finally, as a consequence of the described build system improvements, the CI configuration has been simplified as fewer build combinations need to be tested, which in turn has enabled us to integrate additional build jobs into the CI, such as automated Python wheel builds, Windows installer builds, test code coverage and documentation generation.

Thanks to having centralized documentation and automated documentation building in the CI, developers are now able to more easily find information on how to build, install, configure, debug, profile, measure test code coverage, manage releases and versioning, and build Python wheels. The user documentation has also become more accessible and searchable, which makes NEURON more accessible to the community.

Since the introduction of test code coverage tracking we were able to increase test coverage by almost 18%. For the code components that are being maintained and developed by the NEURON community, more than half the code is covered by tests. Introducing systematic testing and coverage reports has allowed us to keep track of our progress and facilitate the refactoring and maintenance of the code. Up to date code coverage reports are available at https://app.codecov.io/gh/neuronsimulator/nrn.

### 3.2 Improved software and hardware portability

#### 3.2.1 streamlined NEURON software Distributions

Building distributions (Windows and macOS installers, Debian packages) and testing them reliably on different platforms against different software toolchains has historically been a major hurdle to NEURON releases. This had been extending release cycles and delaying the introduction of new features. With the CI pipelines described in Section 2.1.2 and a modernized CMake-based build system, we are now able to automatically build portable NEURON Python wheels for Linux and macOS (including Apple M1) and an installer for Windows. With the infrastructure in place we are able to offer both nightly wheels and installers, containing the latest changes, as well as more rigorously tested, and hence stable, release builds. By simplifying, automating and documenting the build steps, we have streamlined the process of creating new NEURON releases.

Python’s dominance in scientific workflows and the widespread use of Python-based data processing, analysis and visualization tools in the scientific community can be attributed to the ease-of-use and portability of these packages. By distributing NEURON as a Python package we are embracing this trend and making it easier to adopt NEURON. These Python wheels, as well as binary installers, provide a full, portable distribution of NEURON targeting desktop environments, cloud instances and HPC clusters alike. Although we do not currently provide a Python wheel for Windows, users can make use of the Windows Installer or the Linux Python wheel using Windows Subsystem for Linux (WSL). All these distributions have support for dynamically loading MPI, Python, and Interviews graphics. This ease of use has made distribution via Python wheels the preferred way of installing NEURON. For Python wheels, we currently see an average of around 3000 downloads/month, and over 17’000 downloads in the last six months. Released wheels are available via https://pypi.org/project/NEURON.

#### 3.2.2 Improved hardware portability

Supporting a wide range of use-cases requires strong hardware support for architectures ranging from laptops to cloud and HPC platforms. Thanks to its updated and improved build system it is straightforward to build NEURON and CoreNEURON for a variety of hardware platforms including x86, ARM64, POWER and NVIDIA PTX. The respective vendor compilers are able to take advantage of CoreNEURON’s improved data-structures and produce optimized code. In order to make CoreNEURON’s GPU backend accessible to the wider user community we have additionally created NEURON Python wheels with GPU acceleration enabled. Currently these specialized wheels can only be used on environments with NVIDIA GPUs with compute capability 6, 7 or 8 and the NVIDIA HPC SDK version 22.1 with CUDA 11.5. These wheels can be downloaded from https://pypi.org/project/NEURON-gpu.

### 3.3 Performance improvements through tighter integration

As presented in Section 2.2, we have greatly improved the integration between NEURON, CoreNEURON and NMODL both on the level of the code organization and their ease of use at runtime. CoreNEURON and NMODL are now git submodules of the nrn repository, allowing us to build single software distribution packages containing optimized CPU and GPU support via CoreNEURON. This allows the user to transparently take advantage of modern hardware platforms such as GPUS, and recent hardware features such as AVX-512 on Intel CPUs. To this end, CoreNEURON’s GPU implementation has been made production-ready, allowing easy offloading to NVIDIA GPUs. More importantly, the newly introduced in-memory transfer mode allows CoreNEURON simulations to be directly called from NEURON and the model state to be passed back and forth between NEURON and CoreNEURON. The workflow to utilize these new features is illustrated in Figure 6.

**Figure 6:**
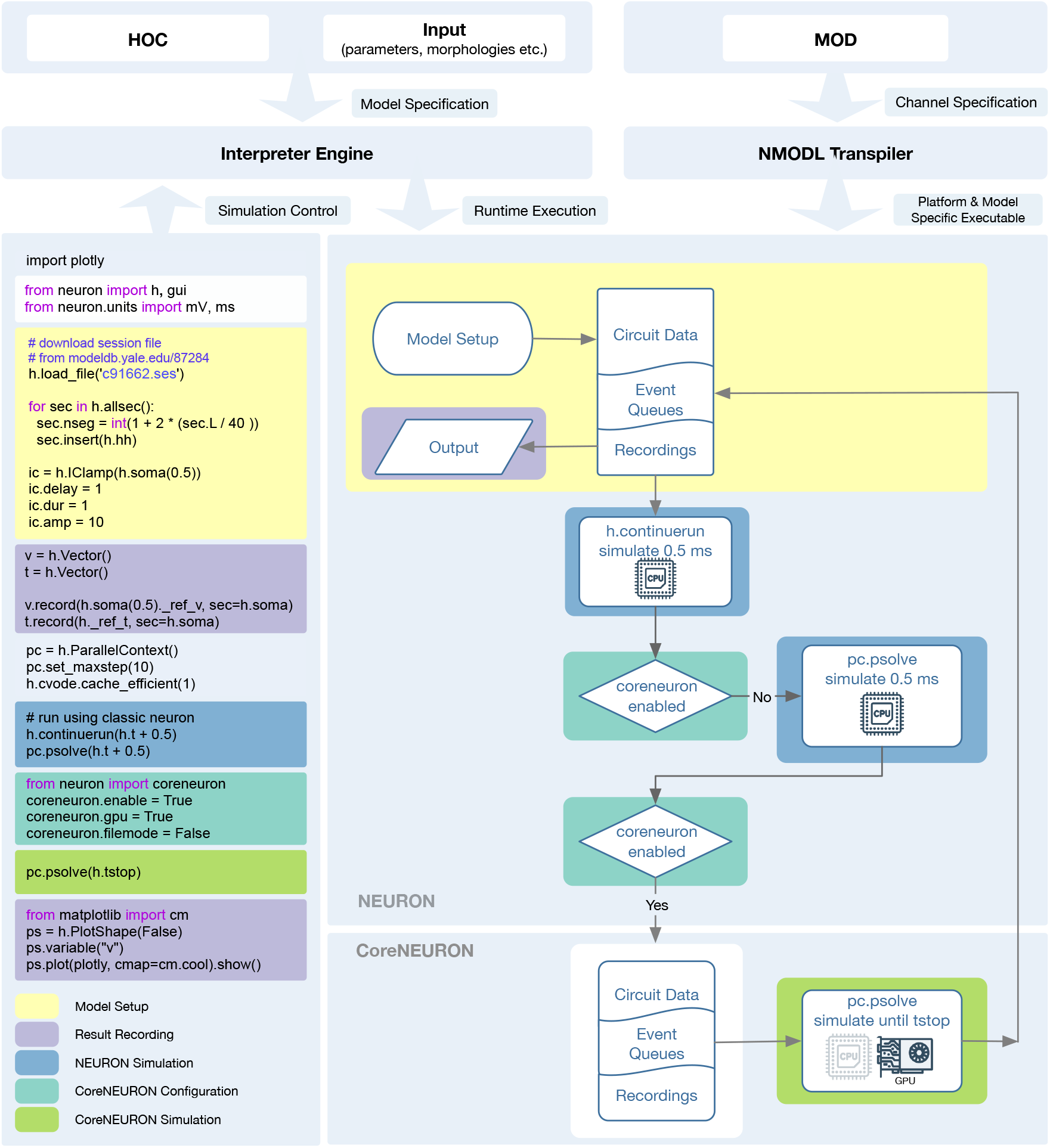
Diagram showing the NEURON and CoreNEURON execution workflow using the Python API: The Python code on the left demonstrates the CoreNEURON Python API usage and interoperability between NEURON and CoreNEURON solvers. The different shaded areas in the code correspond to the boxes of the same color on the right. First, the script sets up the model in NEURON and defines the entities to be recorded. In the following `h.continuerun()` statement the script starts by running the simulation in NEURON for 0.5 ms. Since the `coreneuron` and `gpu` options have not been enabled yet, the following call to `pc.psolve()` advances the simulation for 0.5 ms using NEURON. The call to `pc.psolve()` that is executed after enabling CoreNEURON, however, first copies the model to CoreNEURON using the direct-mode transfer and then runs the simulation until the prescribed `h.tstop` using CoreNEURON on GPUs. After finishing the CoreNEURON simulation step all variables and events are transferred back to NEURON.

After constructing and initializing the model using NEURON’s session file, simulation is first run in NEURON with `h.continuerun()` and `pc.psolve()`. GPU support via CoreNEURON is enabled using the newly introduced `coreneuron` Python module. Having CoreNEURON enabled will cause `pc.psolve` to use the CoreNEURON solver instead of the default NEURON solver. All necessary model state is automatically transferred to CoreNEURON before the simulation is continued using the CoreNEURON solver. At the end of the solver call, the model state is transferred back to NEURON, allowing user to save necessary recording results.

With this tighter integration, it is now possible to easily switch a large number of models, notably those using the fixed timestep method, to use the optimized CoreNEURON solver. To quantify the performance benefits, in this section we showcase three different models that were ported to use CoreNEURON. We will compare the performance of NEURON and CoreNEURON running on CPU and GPU on the olfactory 3D bulb model 3.3.1, the rat CA1 hippocampus model 3.3.2 and the rodent motor (M1) cortical model 3.3.3.

The benchmarking system, Blue Brain 5 (Phase 2), with its hardware and software toolchains is summarized in Table 1. This system is based on an HPE SGI 8600 platform (HPE, 2022) and is housed at the Swiss National Supercomputing Center (CSCS). The GPU partition of the system has compute nodes with Intel Cascade Lake processors and NVIDIA Volta 100 GPUs. We used vendor-provided compiler toolchains and MPI libraries for the benchmarking. Unless otherwise specified, measurements were performed using two compute nodes, providing a total of 80 physical cores. We ran all CPU benchmarks in pure MPI mode by pinning one MPI rank per core. For GPU executions we reduced the number of MPI ranks to 16 (eight ranks per node) in order to achieve better utilization, enabled the CUDA Multi-Process Service (MPS) and used two or four NVIDIA V100 GPUs on each node. For the CPU measurements, NEURON and CoreNEURON were compiled using the Intel C++ compiler, while for GPU measurements CoreNEURON was compiled using the NVIDIA C++ compiler. All CoreNEURON benchmarks were both performed using the legacy MOD2C transpiler and the next-generation NMODL transpiler. All reported speedups were averaged over ten runs.

**Table 1:**
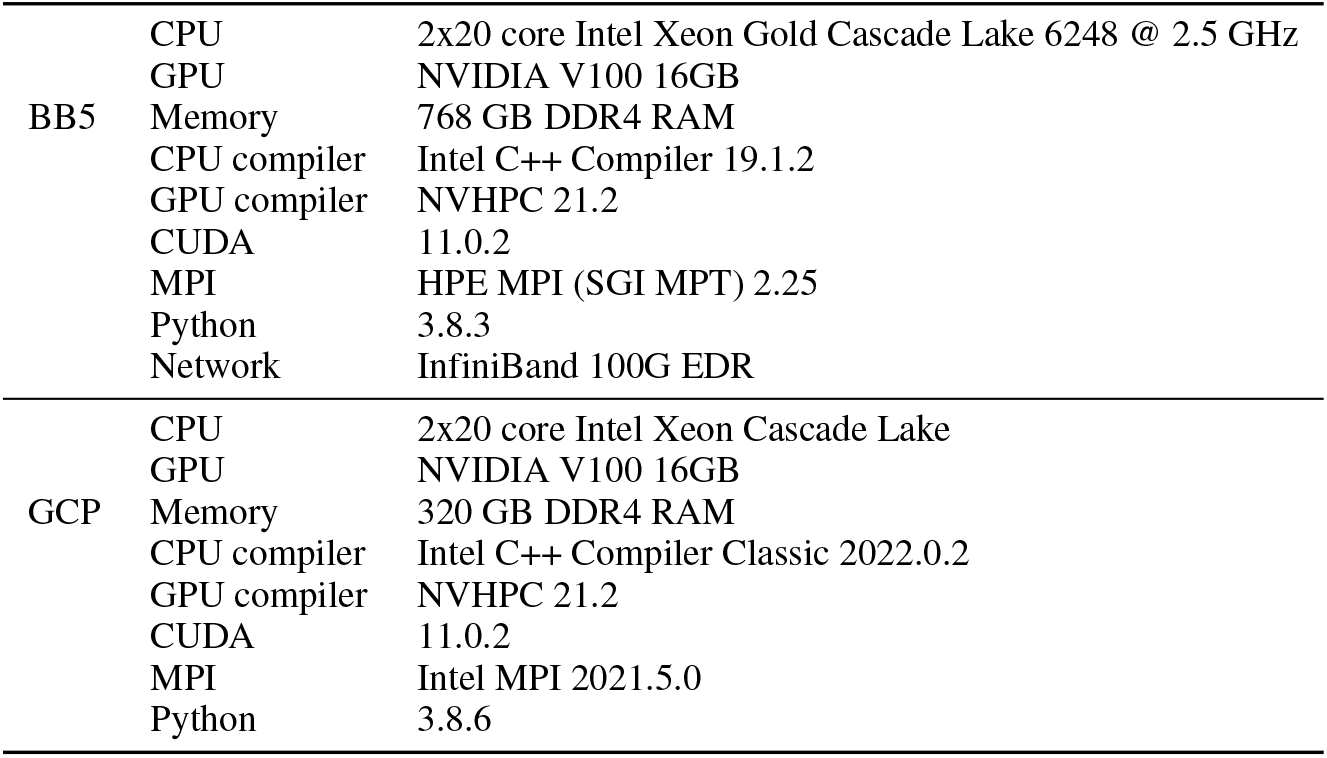
Details of Benchmarking Systems: Blue Brain 5 (Phase 2) Supercomputer and Google Cloud Platform

#### 3.3.1 Accelerating 3D Olfactory bulb model simulations via CoreNEURON

The olfactory bulb microcircuit developed by Migliore et al. (2014), serves as a model for studying the functional consequences of the laminar organization observed in cortical systems. The model was developed using realistic three-dimensional inputs, cell morphologies and network connectivity. The original model uses Python 2 and is available on ModelDB; we updated this model to use Python 3 and CoreNEURON to run our benchmarks. The updated version is publicly available from the GitHub repository of the Human Brain Project https://github.com/HumanBrainProject/olfactory-bulb-3d. The full model consists of 191,410 cells, 3,388,282 synapses and a total of 9,118,745 compartments. We simulated the default model configuration with a biological duration of 1050 ms and a timestep of 46.875 μs.

Figure 7 shows the performance difference between NEURON and CoreNEURON solvers when executing on CPU and GPU hardware. As can be observed in Figure 7-A, the simulation using CoreNEURON on two full CPU nodes is 3.5 × faster than the baseline NEURON benchmark run on the same hardware. The achieved acceleration is due to the use of SIMD instructions, which is enabled by the efficient internal data structures and the SoA (Structure of Array) memory layout used by CoreNEURON. When GPU offloading is enabled in CoreNEURON then with two GPUs per node the speedup increases to 21.4 × compared with the baseline NEURON benchmark. Performance does not scale linearly when doubling the number of GPUs per node to four, and we see a maximum speedup of 30.4×. This is due to more time spent in the communication between the eight GPUs and the size of the model reaching the strong-scaling limit. In order to understand the performance differences between CoreNEURON executing on CPU and GPU Figure 7-B shows a comparison of the two runtime profiles broken down into the most relevant execution regions. We have normalized the time of each region with the total execution time of the simulation. On the one hand, Figure 7-B shows that the relative time spent in the most compute intensive parts such as the `Current calculation` and the `State update` is reduced significantly when executing on the GPU. The Hines `Matrix solver` does not currently benefit from GPU acceleration. This is due to data dependencies and limited parallelism inherent to the algorithm. On the other hand, we can see that the event delivery and CPU-GPU data transfer incur an additional cost compared to the CPU execution. Also, the spike exchange routines take a larger share in runtime on the GPU than on the CPU. This shows that the highly parallelizable compute operations are readily accelerated on the GPU while data movement and code with higher execution divergence are favored by the CPU.

**Figure 7:**
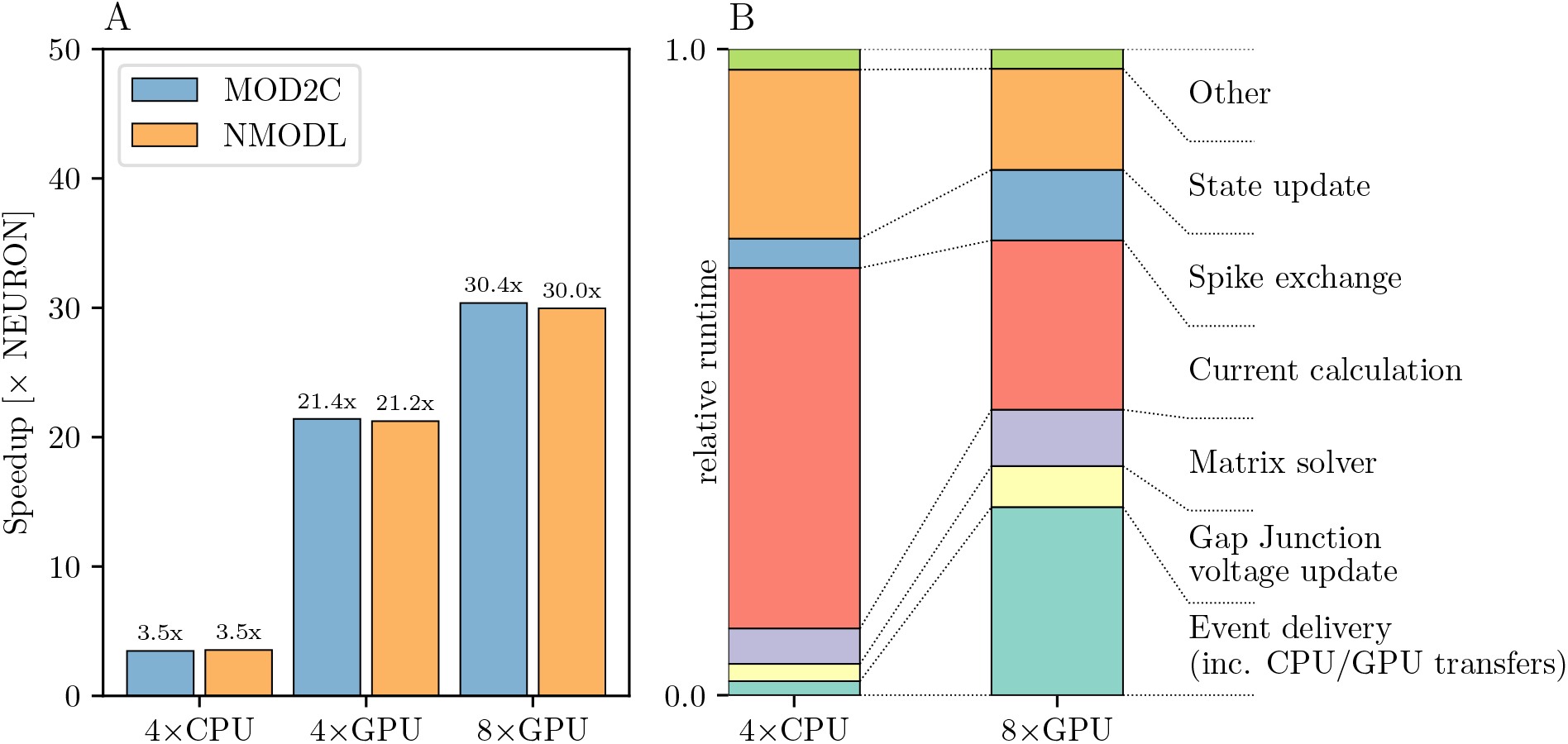
Olfactory 3D bulb model performance comparison: **(A)** improvement in the simulation time using CoreNEURON on CPU and GPU, with respect to NEURON running on CPU. We show the speedups using both MOD2C and NMODL transpilers. **(B)** comparison of the two runtime profiles of the CoreNEURON run on CPU and GPU. The relative time (normalized to the total execution time) of the different execution regions in one timestep is shown. All benchmarks were run on two compute nodes with a total of 80 MPI ranks.

#### 3.3.2 Accelerating Rat cA1 Hippocampus simulations using GPUs

Another interesting example of a NEURON based simulation is the full-scale model of the rat hippocampus CA1 region built as part of the European Human Brain Project (manuscript in preparation). A recent draft of this model contains 456,378 neurons with 12 morphological types and 16 morpho-electrical types. The CA1 neurons in this model employ up to 11 active conductance classes, with up to 9 of those classes used in the dendrites. This model is used for running various large scale in-silico experiments on different European HPC systems. A reduced but representative version of this model is used for and available as part of the Hippocampus Microcircuit Massive Open Online Course^7^ offered on edx.org.

Simulating the full rat CA1 hippocampus model is computationally expensive and requires large-scale HPC systems, due to its scale and biologically detailed nature. It represents, therefore, an ideal showcase for the improvements in hardware portability described above, most notably the GPU support provided by CoreNEURON. For the purpose of this showcase, we used aforementioned reduced model of the CA1 hippocampus containing 18,198 cells with 11,401,920 compartments and 107,237,482 synapses. Furthermore, we simulated a biological duration of 1000 ms with a timestep of 25 μs. Finally, the runtime configuration for the GPU benchmarks had to be adjusted so that 80 (instead of 16) MPI ranks were used. This was necessary due to a technical limit on the number of artificial cells of a given type that CoreNEURON can simulate in a single MPI rank.

Figure 8-A shows that using CoreNEURON yields a performance improvement of 3 - 4× compared with NEURON when executing on four Cascade Lake CPUs. When enabling GPU offloading in CoreNEURON it is possible to achieve up to 52 × improvement compared with the NEURON baseline. It is worth mentioning that the new NMODL transpiler shows significant improvement in execution time over the legacy MOD2C transpiler (i.e. a speedup of 52 × vs 42 × when using eight GPUs). This is due to NMODL’s analytic solver generation, which is based on SymPy (Meurer et al., 2017) and the Eigen library (Guennebaud et al., 2010). The performance improvement can be mainly attributed to the fact that `State update` kernels represent the evaluation of `DERIVATIVE` blocks in the `MOD` files, which contain ODEs that are now efficiently solved by NMODL. It is apparent that due to the computationally expensive nature of this model it scales linearly from four to eight GPUs and executing on GPUs provides a substantial benefit over the baseline NEURON benchmark. Analogous to Figure 7-B, Figure 8-B shows the relative breakdown of the total execution time for the most relevant regions. In contrast to the previous example, however, we see here that the current calculation and state update remain dominant on the GPU, while event delivery and CPU-GPU transfer play a smaller role. We interpret this result as an indication that this simulation is better suited for the strengths of the GPU hardware than the olfactory bulb model.

**Figure 8:**
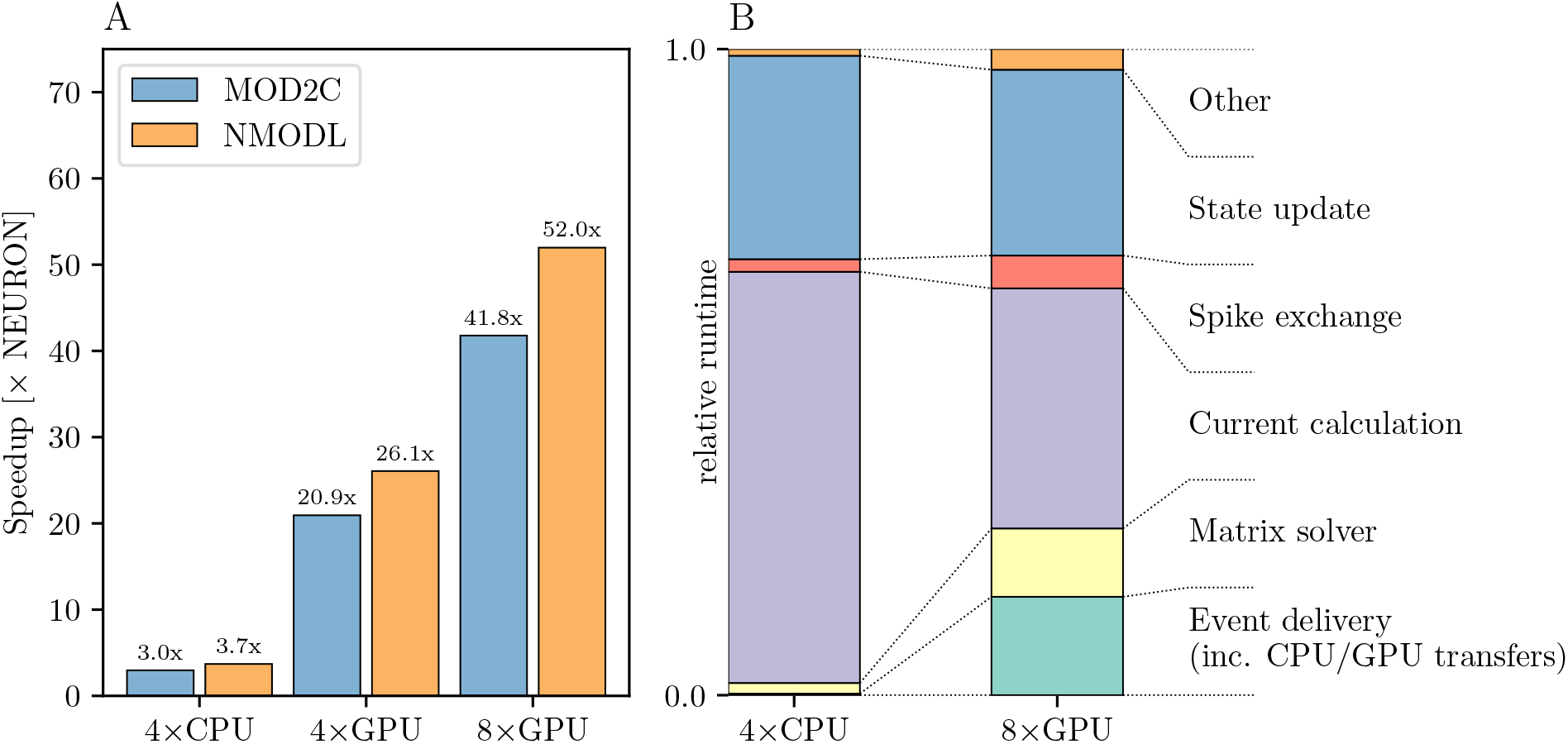
Rat CA1 hippocampus model performance comparison: **(A)** improvement in the simulation time using CoreNEURON on CPU and GPU, with respect to NEURON running on CPU. NMODL shows a significant improvement compared to the legacy MOD2C transpiler, which is due to analytic solver support in NMODL. **(B)** comparison of the two runtime profiles of the CoreNEURON run on CPU and GPU. The relative time (normalized with the total execution time) of the different execution regions in one timestep. All benchmarks were run on two compute nodes with a total of 80 MPI ranks.

#### 3.3.3 Simulating large-scale cortical models with NetPyNE

Here we report on the integration of NetPyNE with the CoreNEURON solver, thus taking full advantage of modern CPU/GPU architectures to reduce the simulation time. To illustrate this, we compared the performance of a large-scale and biologically realistic model of mouse motor (M1) cortical circuit on Google Cloud Platform (GCP) as well as Blue Brain 5. The mouse M1 model simulates a 300 μm cylinder of cortical tissue including 10,651 neurons, of three excitatory and four inhibitory types, and 38 million synapses, with 7,000 spike generators (NetStims) simulating long-range inputs from seven thalamic and cortical regions. All model neuronal densities, classes, morphologies, biophysics, and connectivity were derived from experimental data. The model is being used to study neural coding, learning mechanisms and brain disease in motor cortex circuits (Dura-Bernal et al., 2022b; Sivagnanam et al., 2020).

GCP offers a wide variety of machine types tailored to different applications. For our benchmarks we allocated `n2-standard-80` and `nvidia-tesla-v100` nodes. The `n2-standard-80` are dual-socket Intel Cascade Lake based nodes while `nvidia-tesla-v100` compute nodes consist of dual-socket Cascade Lake based systems with 8 NVIDIA V100 GPUs. NEURON and CoreNEURON CPU measurements of the M1 NetPyNE benchmarks were run using 80 MPI ranks evenly distributed over two `n2-standard-80` nodes, while GPU runs were performed with 16 MPI ranks on one node pinning one MPI rank per core and distributing the ranks evenly between 4 or 8 GPUs. To allow for a fairer comparison between on-premise HPC hardware and cloud platforms we adjusted the benchmark configuration accordingly on Blue Brain 5 running with 16 MPI ranks for the GPU runs while maintaining the 80 ranks for the CPU runs. Figure 9-A shows benchmark performance on Blue Brain 5. The CoreNEURON solver is 4× faster compared to NEURON when executing on CPU only. When GPU support is enabled then we achieve speedup of 26 × and 39 × with four and eight GPUs respectively. Similarly to Section 3.3.1 the suboptimal scaling from four to eight GPUs suggests that the model’s size and computational intensity do not fully saturate the allocated hardware and do not fully benefit from it. Figure 9-B shows the performance improvements achieved on GCP. The measured speedups compared with the baseline NEURON CPU runs are 3.6×, 21 × and 30× for CoreNEURON on CPU, on four and eight GPUs respectively. While the achieved speedups on GCP are slightly lower than on Blue Brain 5, they still show a clear performance advantage when using CoreNEURON with GPU support. Furthermore, this shows that one can get improved performance in cloud environments just as on traditional, on-premise cluster systems. Finally, we note that the use of NEURON Python wheels with their built-in GPU support drastically simplifies the efforts to setup the NEURON simulator toolchain with GPU support in cloud environments.

**Figure 9:**
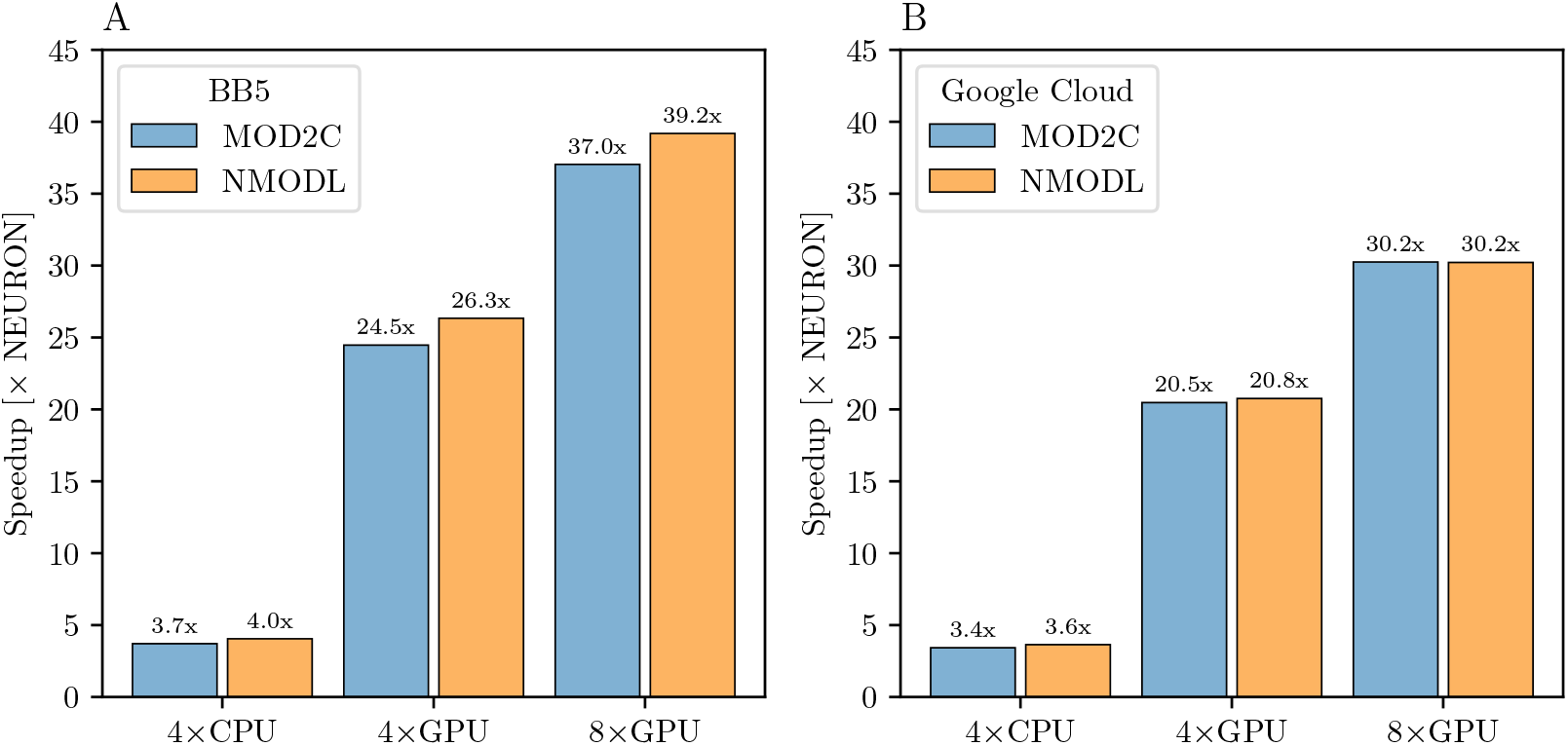
M1 cortical model performance comparison: **(A)** improvement in simulation time running CoreNEURON on CPU and GPU on the Blue Brain 5 supercomputer with respect to NEURON running on CPU. CPU benchmarks were run with 80 MPI ranks evenly distributed over two nodes, one rank per core. GPU benchmarks used 16 MPI ranks evenly distributed over two nodes. **(B)** improvement in simulation time running CoreNEURON on CPU and GPU on Google Cloud Platform compute resources. The same configuration was used, except for GPU benchmarks being executed on one node.

#### 3.3.4 Improvements in RxD Performance

The developments described in Section 2.4 substantially improved the performance and scalability of reaction-diffusion simulations in NEURON both for 1D and 3D simulation. Using the same model code to simulate pure diffusion in a 1D test cylinder of length 200 μm and diameter 2 μm, with 1001 segments runs 4.2× faster in NEURON 8.0 than in NEURON 7.6.7. We note that “1D” is a slight misnomer here, as NEURON’s version of 1D simulation uses the Hines algorithm (Hines, 1984) which provides 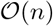 scaling for implicit simulation and supports an arbitrary tree-like morphology where multiple 1D sections may connect at the same point. Likewise, a NEURON implementation of the circadian model of Leloup et al. (1999), a 10-species model with 21 reactions and no diffusion, ran 17.3× faster in NEURON 8.0 than in 7.6.7 (Figure 10). In the NEURON tutorial, we provide a version of this model that requires recent versions of NEURON due to its use of the `neuron.units` submodule to cleanly specify units appropriate for this model, however the ModelDB entry for this paper provides an equivalent representation that runs in both older and recent versions of NEURON.

**Figure 10:**
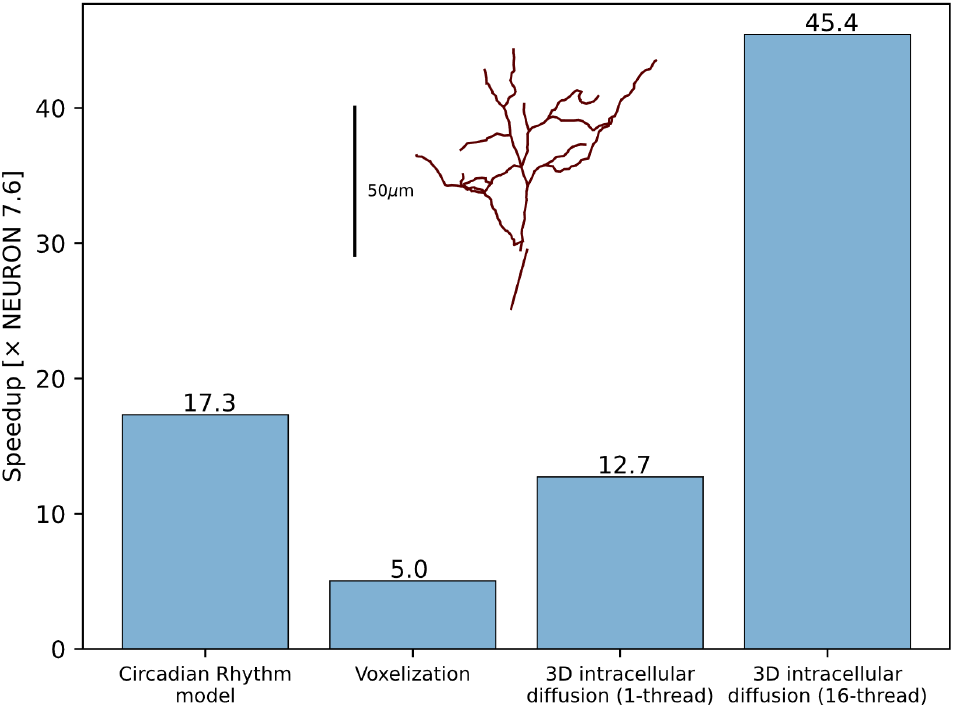
RxD performance improvements for the Circadian Rhythm reaction model with just-in-time compilation, 3D morphology voxelization and intracellular diffusion using the DG-ADI method. The speedup was measured between NEURON 7.6.7 and NEURON 8.0. Inset shows the morphology used for voxelization and 3D diffusion.

Thanks to the improved voxelization method, the construction of voxel representations is now significantly faster. For example, the morphology (with voxel size 0.25 μm in each dimension) of the soma and apical dendrite of a rat CA1 pyramidal neuron (Malik et al., 2016; Ascoli et al., 2007) within 70 μm of the soma completed in 69 seconds in NEURON 8.0 and 349 seconds in NEURON 7.6.7, approximately a 5-fold speedup. The relative volume error (computed by comparing to the volume calculated as above vs with voxel size 0.05 μm in each dimension) was 0.15 % in NEURON 8.0 vs 16 % in NEURON 7.6.7, a reduction made possibly by switching from including the full volume of boundary voxels to a subsampled fractional volume.

3D simulation times are likewise reduced in NEURON 8.0, through the combined effects of eliminating Python from the simulation-time code, compilation of reaction specifications, and the replacement of the simulation algorithm (7.6.7 used SciPy’s (Virtanen et al., 2020) Biconjugate Gradient Stabilized iterative solver for matrix inversion; 8.0 uses a threaded Douglas-Gunn Alternating Direction Implicit method). Pure diffusion over 1 second of simulated time on the cell volume described above with initial conditions of 1 mM in a section of the apical dendrites and 0 mM elsewhere required 18,349 seconds of real time in NEURON 7.6.7, 1,442 seconds of real time in NEURON 8.0 with 1 thread (a 12.7× speedup), and only 426 seconds in NEURON 8.0 when using 16 threads (a 45.4× speedup). (We specify the initial conditions here as the iterative solver used in 7.6.7 would potentially require a different number of iterations depending on the initial conditions; the solver used in 8.0 performs the same calculations each time regardless of the concentrations.)

## 4 Discussion

### 4.1 Sustainability of the NEURON Simulator

Over the years, the scientific community has used NEURON to study a wide range of biophysical questions across various spatial and temporal scales, with over 2500 publications reporting such usage ^8^. More than 750 published NEURON model source codes are available through SenseLab’s ModelDB repository (McDougal et al., 2017), which provides curated metadata describing the biological assumptions. It is fair to say that for biophysically detailed models of neurons and networks, NEURON has become the de facto standard in the community. In light of this, it is quite obvious that continuity of the NEURON project is of great importance to the community.

However, for the majority of its more than 35 years of history, NEURON has been developed under what Gewaltig and Cannon (2014) dubbed a “heroic development model”. While open source from its inception, until recently, the majority of contributions to the NEURON software came from its original author, Michael Hines, and a small group of collaborators from Duke and Yale University. We have now established a radically updated development model and life cycle of the NEURON software towards a modern and collaborative process, facilitating the contribution of modifications from other developers. In the language of Gewaltig and Cannon (2014) we were able to evolve NEURON’s “heroic development model” to a “collaborative development model” with community engagement.

Sustainability of a software project, or its continued development, is of course not a goal in and of itself. A software with the history of NEURON should maintain as much backward compatibility with previously published models as possible, while catering for new use cases. In this respect, we demonstrated how it is possible to process the widely used NMODL language with a modern, compiler-based approach capable of producing highly optimized code, while keeping maximal backward compatibility.

### 4.2 From desktop to supercomputers

For first time users, NEURON is often introduced through interactive explorations on a desktop or laptop, combining small bits of Python with NEURON’s built-in graphical user interface (GUI). This usage remains common in practice, with roughly one-third of NEURON models on ModelDB (McDougal et al., 2017) containing a `.ses` file generated from NEURON’s GUI. Our improved continuous integration and build automation caters to this demand and produces pre-compiled binaries for the most common consumer computing architectures (Windows, macOS, Linux), which allows users to easily install NEURON, including its graphical interface, on their personal systems.

With increasing use, it is common to complement or replace GUI-based use with the power and flexibility of scripting. This allows for easy handling of more diverse models and settings, parameter sweeps, or the evaluation of model variants to establish distributions and robustness of results. Over the last decade, Python has become the language of choice in the practice of computational science, and neurosimulations have been no different (Muller et al., 2015). Since this early integration, both neurosimulators and Python, have evolved substantially, as has their usage pattern. This requires that we express the functionality of NEURON in a Pythonic style and enable simple installation of platform-specific NEURON modules. Here, the adoption of modern Python wheels has dramatically simplified the use of NEURON and user-specific extensions in the Python environment.

As models and simulations become bigger, it is important that the simulator can optimally use the available hardware. In these cases it is particularly useful to be able to rely on NEURON binaries tailored to the computer architecture. This can be achieved through specialized pre-built NEURON binaries or through compiling NEURON with platform-specific parameters, including the support of the CPU’s vector instructions, multiple threads or the use of GPUs (see also Section 4.4).

While there is already a wide range of computer architectures in use today, it is to be expected that the diversity will further increase. This general trend is driven by the challenges of further miniaturizing transistors and performance gains of future computer architectures in part will have to come from specialization (Hennessy and Patterson, 2019), already prominently visible in the field of machine learning (Reuther et al., 2019). Not all of these specializations will necessarily be useful for neurosimulations, but the better we understand the computational costs of our models in neuroscience (Cremonesi et al., 2020; Cremonesi and Schürmann, 2020), the more we may be interested in adopting some of those platforms for specific applications. Here, our work on translating the computationally intensive parts of a neuron model described in the NMODL language into source code that can be compiled for a wide range of computer hardware, in combination with the reduced memory footprint of CoreNEURON, is a major step forward to leverage these developments in the computer industry for neuroscience purposes.

With pre-compiled binaries for all major operating systems (Windows, macOS, Linux), Python scriptability, and built-in support for serial, threaded, MPI, and GPU accelerated calculations, NEURON can readily be used in many computing environments. The new NMODL framework furthermore can be extended to support future computer architectures without having to compromise performance on other platforms.

These efforts for efficiently using today’s computer architectures, are complemented with NEURON’s ability to run on large number of networked nodes, so-called clusters. In previously published work (Hines et al., 2011a), NEURON has run simulations with up to 128,000 processes catering even to the largest models. Nowadays, many universities maintain their own high performance computing environments for such purposes and a growing number of e-infrastructure providers offer high performance computing. For example, smaller numbers of computing resources are freely available for neuroscience simulation with NEURON and other simulators through a Neuroscience Gateway account (Sivagnanam et al., 2013). The EBRAINS research infrastructure provides large supercomputer allocations with preinstalled NEURON and even models for approved research projects.

### 4.3 NEURON as a building block for scientific workflows

Cloud-based Jupyter notebook providers have recently become another accessible way to use NEURON. EBRAINS, developed through the Human Brain Project, provides a Jupyter notebook cloud environment with NEURON and other simulator software pre-installed. Additionally, many public Jupyter servers, including Google Colab, allow installation of Python packages including NEURON via `pip`. Using NEURON through a cloud-based Jupyter server makes it accessible through any computing device with a modern browser, including phones and tablets, and facilitates sharing and collaborating on whole models and examples. To increase NEURON’s usability in Jupyter notebooks, we have added built-in support for Python graphics (including via Plotly and Matplotlib) to NEURON’s `ShapePlot` and `RangeVarPlot` classes, which provide, respectively, a 3D false-color view of the cell and a plot of state values along a path.

Building on top of the modular structure and APIs of NEURON, there are various community tools that have incorporated NEURON as a building block to develop higher level tools. For example, the Human Neocortical Neurosolver (Neymotin et al., 2020) provides its own graphical interface and launches NEURON in a separate process to simulate the underlying neocortical model. NetPyNE’s online graphical interface launches virtual machines with NEURON on demand in the cloud and uses NEURON or CoreNEURON to run simulations. The example of the M1 model demonstrates that in this setting we observe performance increases when switching the simulator from NEURON to CoreNEURON (4× on a tightly coupled cluster, and 3.6 × on a cloud instance). Also, we see good scaling when using multiple GPUs both for the tightly coupled cluster and the cloud; see the next section for a more detailed analysis. These results show that we can tap into the performance improvements of NEURON when invoking it from other tools. Thus, the same improvements in packaging and platform specific optimization that were demonstrated above will benefit also other tools, such as LFPy (Lindén et al., 2014) and BluePyOpt (Van Geit et al., 2016), which rely on NEURON’s simulation capability.

### 4.4 Increased performance for tackling new scientific questions

In recent years, several large-scale and biophysically detailed models have been developed (Markram et al., 2015; Billeh et al., 2020; Casali et al., 2019; Hjorth et al., 2020; Dura-Bernal et al., 2022a,b; Borges et al., 2022). Some of these models are no longer only models of networks and their signal processing but are actually biophysical models of the tissue itself (Markram et al., 2015), capturing a diverse set of properties of the modeled brain region, also referred to as “digital twins”. Such models have proved useful to bridge anatomy and physiology across multiple spatial and time scales (Reimann et al., 2017; Newton et al., 2021).

These types of models have become possible because of more quantitative data, new computational approaches to predicting and inferring missing parameters, and the increase in computer performance. In fact, a large number of improvements to the NEURON software over the last decade were motivated by these types of models: a major common theme in these developments was functionality to support parallel execution on multiple compute nodes (Migliore et al., 2006; Hines et al., 2008a,b, 2011b; Lytton et al., 2016), complemented by platform-specific optimizations (Ewart et al., 2015; Kumbhar et al., 2016). In particular, the platform-specific optimizations underwent a disturbing trend: optimizations of NEURON for the first vector computers had been discontinued in favor of memory optimizations for out-of-order CPUs with intricate cache hierarchies, only to return to SIMD analogous structure-of-array memory layouts for today’s CPUs/GPUs with wide vector units. Previously, this led to special code versions with significant development efforts to adapt the code for different generations of hardware platforms and programming models.

Our work on CoreNEURON (Kumbhar et al., 2019) and NMODL (Kumbhar et al., 2020) has made it possible to contain the platform specific optimizations in the code generation framework, requiring less platform specific code in NEURON and allowing models to remain comparatively free of platform-specific details. Here, we introduced two recent advances. First, NMODL and CoreNEURON are now able to automatically generate code not only for CPUs but also GPUs. Second, the transparent integration with NEURON makes it possible to leverage this capability simply as an accelerator for simulations on a user’s desktop, or to dramatically speed up large-scale simulations on supercomputers.

We compared running three different large scale models with NEURON and CoreNEURON in different hardware configurations. Compared to the baseline running with NEURON, transparently offloading to CoreNEURON achieves a four-fold speed-up on the same CPU hardware. This performance increase is mostly due to the better utilization of the vector units, a more cache efficient memory layout, and less data transfer between the CPU and main memory (Kumbhar et al., 2019). We furthermore demonstrate that it is now also possible to seamlessly make use of NVIDIA GPU hardware. We demonstrated a speed-up of 30×, 39× and 52× when using eight GPUs for the olfactory bulb model, the cortical M1 model and the hippocampus CA1 model, respectively, compared to four full CPUs. For the hippocampus CA1 model, we see ideal scaling up to 52× when doubling the GPU number. Both the thalamocortical M1 model and the olfactory bulb model show suboptimal scaling when moving from four to eight GPUs (39× and 30×, respectively), which we attribute to their lower computational cost compared with the hippocampus CA1 model, leading to lower utilization of the GPUs, and an overall lower compute-to-communicate ratio seen in the relatively longer time spent in event delivery and CPU-GPU data transfers.

These numbers should not be used for comparing the CPU’s suitability for neurosimulations with that of a GPU. As the two architectures have wildly different characteristics in terms of total floating point performance and memory bandwidth, a deeper analysis is required to establish the efficiency of the simulations on the respective platforms (Lee et al., 2010). However, what can clearly be demonstrated with these numbers is that it is now possible for any user to readily make use of NVIDIA GPUs if they are installed.

### 4.5 Simulations in the cloud

Low-cost virtual machines or dedicated servers are now also available from commercial providers billed by the second - typically referred to as “the cloud”. Using NEURON in these environments requires that one can quickly configure NEURON there. In these environments, it is furthermore desirable to save intermediate results in a database to allow examining the results mid-calculation and to facilitate resuming in the event of timeouts and other issues, which can be readily done with database functionality integrated in Python (such as SQLite3). This is where our work on a straightforward `pip install` for NEURON is particularly useful, as more generally described in Sivagnanam et al. (2020) as a practical approach for both small sets of simulations and very large ones. Not only could we demonstrate the feasibility of running in the cloud using the Google Compute Cloud and NetPyNE with CoreNEURON as the compute backend, but we could also demonstrate that the achievable performance on a moderate set of nodes (moderate when compared to clusters and supercomputers) is on par with that of bare metal simulations on dedicated cluster. Using CoreNEURON showed a speedup that is comparable (3.6× vs. 4.0×) with the one achieved on a dedicated cluster. Using GPU offloading with CoreNEURON offered a 30× speedup compared with the baseline NEURON simulation. These results should be put in the context of simple on-demand access to such compute resources, NEURON’s integration in an ecosystem of computational neuroscience, data processing and analysis tools.

### 4.6 Efficiently integrating subcellular and extracellular detail into neurosimulations

Some of the greatest health challenges of our time like stroke (the #2 cause of death worldwide) and dementia (#7 cause of death worldwide) are driven by the interaction of effects spanning from the sub-cellular level to neuron, network, and organism levels. These scales have often been addressed separately using different techniques with computational neuroscientists focusing on the neuron and network level while systems biologists study the protein interactions and systems-level questions, but this split has long been viewed as artificial and possibly problematic (see e.g. De Schutter (2008)). Exporting point dynamics to SBML for exploration with systems biology tools is an explicit bridge across this divide, but we believe that within NEURON the recent enhancements to NEURON’s reaction-diffusion (RxD) support provide a conceptual bridge, supporting studies that were previously impractical. Faster simulations in both 1D and 3D are more than just a convenience for the modeler: they allow more detailed dynamics to be simulated in the same amount of time featuring a more complete representation of molecular interactions. They allow parameter sweeps at a higher resolution of detail, and they allow building more detailed training sets for machine-learning powered approximations of complex biophysics (e.g. Pham et al. (2021)). The recently added ability to use NEURON’s SaveState class with RxD models will facilitate running multiple experiments from a complex initial state and investigating steady state dependence on parameter values. New NEURON features like 3D extracellular simulation allow exploring how detailed cell models interact with each other through the extracellular environment and provide opportunities to include the effects of astrocytes (associated with multiple neurological diseases including Alzheimer’s and Parkinson’s), blood vessels, and other considerations historically under-studied by computational neuroscience.

### 4.7 Outlook

The updated development model, improved build system and enhanced software testing regime presented in this study provide the foundations for further modernization and re-engineering of NEURON. These improvements will enable more complex changes to be made while maintaining correctness and performance. The performance improvements reported here, coupled with the ease-of-use of the newly released Python wheels, show that CoreNEURON could soon become the default simulation engine for NEURON models, and that the improved integration between NEURON and CoreNEURON that we have presented is just the starting point. Further integration will require more migration of NEURON code to C++ and the development of modern data structures that can be exchanged between NEURON and CoreNEURON more easily and efficiently. The next-generation NMODL framework source-to-source compiler is able to parse the entire NMODL language and generate efficient and portable code for most existing `MOD` files. By further extending its code generation capabilities we will be able to replace the legacy NOCMODL and MOD2C DSL translators entirely. Ultimately this will allow the neuroscience community to use NEURON to simulate increasingly complex brain models in more accessible ways on systems ranging from desktops to supercomputers.

## Author Contributions

NTC, WL, MLH and FS conceptualized and led the study. MLH and PK led the overall software development on NEURON, CoreNEURON and NMODL. OA, PK, NC, JGK, OL, IM, FP, AS and MLH, contributed to aspects of NEURON, CoreNEURON and NMODL software development. OL and IM led GPU support integration and performance improvements. AS and NC led software engineering efforts including refactoring, CI and testing. FP implemented support for portable Python wheels for CPU and GPU platforms. IM and SDB performed and validated the NetPyNE benchmarks. IM, OL, PK and OA performed and validated the 3D olfactory bulb and Hippocampus benchmarks. RAM and AJHN led and performed software development on RxD. OA, PK and FS wrote the manuscript. NC, SDB, JGK, OL, IM, RAM, AJHN, FP, AS, TC, WL, MLH contributed to the manuscript. All authors gave feedback to the manuscript.

## Acknowledgements

We thank Alessandro Cattabiani, Christos Kotsalos and Tristan Carel for improving AST visitors and solver support in NMODL. We thank Jorge Blanco Alonso and Christos Kotsalos for improving the reports interface and GPU solver performance in CoreNEURON respectively. We thank Evan Blasy, Lia Eggleston, and Cameron Conte for their respective contributions in SBML export, 3D voxelization, and 3D simulation functionalities in RxD. We thank Michele Migliore for providing the Olfactory bulb model.

## Data Availability Statement

The NEURON simulator, with all the features and improvements described in this paper, is available as version 8.1 in the NEURON GitHub repository^9^. NetPyNE with CoreNEURON support is released as version 1.0.1 and available in the NetPyNE GitHub repository^10^. From the performance benchmarking studies in Section 3.3, the 3D Olfactory bulb model is available in the Human Brain Project GitHub repository^11^, the Rat CA1 Hippocampus model is available as part of the Hippocampus Microcircuit Massive Open Online Course^12^ offered on edx.org, and the M1 cortical circuit is available in the SUNY Downstate Medical Center GitHub repository^13^. All the benchmarking scripts, performance measurement data and figures are available in the NEURON GitHub repository^14^.

## Footnotes

1 https://ebrains.eu/

2 https://www.msys2.org/

3 https://readthedocs.org/

4 http://netpyne.org/models

5 http://gui.netpyne.org

6 Adapted from https://github.com/src-d/hercules

7 https://www.edx.org/course/simulating-a-hippocampus-microcircuit

8 https://www.neuron.yale.edu/neuron/publications/neuron-bibliography

9 https://github.com/neuronsimulator/nrn

10 https://github.com/suny-downstate-medical-center/netpyne

11 https://github.com/HumanBrainProject/olfactory-bulb-3d

12 https://www.edx.org/course/simulating-a-hippocampus-microcircuit

13 https://github.com/suny-downstate-medical-center/M1_NEURON_paper/

14 https://github.com/neuronsimulator/neuron_frontiers_2022_artifacts

## References

Agullo, E., Demmel, J., Dongarra, J., Hadri, B., Kurzak, J., Langou, J., et al. (2009). Numerical linear algebra on emerging architectures: The PLASMA and MAGMA projects. Journal of Physics: Conference Series 180, 012037. doi:10.1088/1742-6596/180/1/012037

Akar, N. A., Cumming, B., Karakasis, V., Küsters, A., Klijn, W., Peyser, A., et al. (2019). Arbor — a morphologically-detailed neural network simulation library for contemporary high-performance computing architectures. In 2019 27th Euromicro International Conference on Parallel, Distributed and Network-Based Processing (PDP). 274–282. doi:10.1109/EMPDP.2019.8671560

Amunts, K., Knoll, A. C., Lippert, T., Pennartz, C. M., Ryvlin, P., Destexhe, A., et al. (2019). The human brain project—synergy between neuroscience, computing, informatics, and brain-inspired technologies. PLoS biology 17, e3000344

Anwar, H., Caby, S., Dura-Bernal, S., D’Onofrio, D., Hasegan, D., Deible, M., et al. (2021). Training a spiking neuronal network model of visual-motor cortex to play a virtual racket-ball game using reinforcement learning

Ascoli, G. A., Donohue, D. E., and Halavi, M. (2007). NeuroMorpho.Org: a central resource for neuronal morphologies. Journal of Neuroscience 27, 9247–9251

Bartlett, R. A., Heroux, M. A., and Willenbring, J. M. (2012). Overview of the TriBITS lifecycle model: A Lean/Agile software lifecycle model for research-based computational science and engineering software. In 2012 IEEE 8th International Conference on E-Science. 1–8. doi:10.1109/eScience.2012.6404448

Beckingsale, D. A., Burmark, J., Hornung, R., Jones, H., Killian, W., Kunen, A. J., et al. (2019). Raja: Portable performance for large-scale scientific applications. In 2019 IEEE/ACM International Workshop on Performance, Portability and Productivity in HPC (P3HPC). 71–81. doi:10.1109/P3HPC49587.2019.00012

Billeh, Y. N., Cai, B., Gratiy, S. L., Dai, K., Iyer, R., Gouwens, N. W., et al. (2020). Systematic Integration of Structural and Functional Data into Multi-scale Models of Mouse Primary Visual Cortex. Neuron 106, 388–403.e18. doi:10.1016/j.neuron.2020.01.040

Blundell, I., Brette, R., Cleland, T. A., Close, T. G., Coca, D., Davison, A. P., et al. (2018). Code Generation in Computational Neuroscience: A Review of Tools and Techniques. Frontiers in Neuroinformatics 12, 68. doi:10.3389/fninf.2018.00068

Borges, F. d. S., Moreira, J. V., Takarabe, L. M., Lytton, W. W., and Dura-Bernal, S. (2022). Large-scale biophysically detailed model of somatosensory thalamocortical circuits in netpyne. bioRxiv doi:10.1101/2022.02.03.479029

Brette, R., Rudolph, M., Carnevale, T., Hines, M., Beeman, D., Bower, J. M., et al. (2007). Simulation of networks of spiking neurons: A review of tools and strategies 23, 349–398. doi:https://doi.org/10.1007/s10827-007-0038-6

Bryson, A., Berkovic, S. F., Petrou, S., and Grayden, D. B. (2021). State transitions through inhibitory interneurons in a cortical network model. PLoS Comput. Biol. 17, e1009521

Carter Edwards, H., Trott, C. R., and Sunderland, D. (2014). Kokkos: Enabling manycore performance portability through polymorphic memory access patterns. Journal of Parallel and Distributed Computing 74, 3202–3216. doi:https://doi.org/10.1016/j.jpdc.2014.07.003. Domain-Specific Languages and High-Level Frameworks for High-Performance Computing

Casali, S., Marenzi, E., Medini, C., Casellato, C., and D’Angelo, E. (2019). Reconstruction and Simulation of a Scaffold Model of the Cerebellar Network. Frontiers in Neuroinformatics 13, 37. doi:10.3389/fninf.2019.00037

Cremonesi, F., Hager, G., Wellein, G., and Schürmann, F. (2020). Analytic Performance Modeling and Analysis of Detailed Neuron Simulations. The International Journal of High Performance Computing Applications 34, 428–449. doi:10.1177/1094342020912528

Cremonesi, F. and Schürmann, F. (2020). Understanding Computational Costs of Cellular-Level Brain Tissue Simulations Through Analytical Performance Models. Neuroinformatics doi:10.1007/s12021-019-09451-w

Crouch, S., Hong, N. C., Hettrick, S., Jackson, M., Pawlik, A., Sufi, S., et al. (2013). The Software Sustainability Institute: Changing Research Software Attitudes and Practices. Computing in Science Engineering 15, 74–80. doi:10.1109/MCSE.2013.133

De Schutter, E. (2008). Why are computational neuroscience and systems biology so separate? PLoS computational biology 4, e1000078

de Verdière, G. C. (2020). Recommendations of the “Extreme Data and Computing Initiative – 2” project, assessment for legacy code and software modernisation. https://exdci.eu/sites/default/files/public/files/d4.5f.pdf. [Online; accessed 14-September-2021

Douglas, J. and Gunn, J. E. (1964). A general formulation of alternating direction methods. Numèrische mathèmatik 6, 428–453

Dura-Bernal, S., Griffith, E. Y., Barczak, A., O’Connell, M. N., McGinnis, T., Schroeder, C. E., et al. (2022a). Data-driven multiscale model of macaque auditory thalamocortical circuits reproduces in vivo dynamics. bioRxiv doi:10.1101/2022.02.03.479036

Dura-Bernal, S., Neymotin, S. A., Suter, B. A., Dacre, J., Schiemann, J., Duguid, I., et al. (2022b). Multiscale model of primary motor cortex circuits reproduces in vivo cell type-specific dynamics associated with behavior. bioRxiv doi:10.1101/2022.02.03.479040

Dura-Bernal, S., Suter, B. A., Gleeson, P., Cantarelli, M., Quintana, A., Rodriguez, F., et al. (2019). Netpyne, a tool for data-driven multiscale modeling of brain circuits. eLife 8, e44494. doi:10.7554/eLife.44494

Einevoll, G. T., Destexhe, A., Diesmann, M., Grün, S., Jirsa, V., de Kamps, M., et al. (2019). The Scientific Case for Brain Simulations. Neuron 102, 735–744. doi:10.1016/j.neuron.2019.03.027

Erdemir, A., Mulugeta, L., Ku, J. P., Drach, A., Horner, M., Morrison, T. M., et al. (2020). Credible practice of modeling and simulation in healthcare: ten rules from a multidisciplinary perspective. J. Transl. Med. 18, 369

Ewart, T., Yates, S., Cremonesi, F., Kumbhar, P., Schürmann, F., and Delalondre, F. (2015). Performance evaluation of the IBM POWER8 architecture to support computational neuroscientific application using morphologically detailed neurons. In Proceedings of the 6th International Workshop on Performance Modeling, Benchmarking, and Simulation of High Performance Computing Systems - PMBS ‘15 (Austin, Texas: ACM Press), 1–11. doi:10.1145/2832087.2832088

Gewaltig, M. and Diesmann, M. (2007). NEST (NEural Simulation Tool). Scholarpedia 2, 1430. doi:10.4249/scholarpedia.1430.Revision#130182

Gewaltig, M.-O. and Cannon, R. (2014). Current Practice in Software Development for Computational Neuroscience and How to Improve It. PLOS Computational Biology 10, e1003376. doi:10.1371/journal.pcbi.1003376

Gleeson, P., Cantarelli, M., Marin, B., Quintana, A., Earnshaw, M., Sadeh, S., et al. (2019). Open source brain: a collaborative resource for visualizing, analyzing, simulating, and developing standardized models of neurons and circuits. Neuron 103, 395–411

Goodman, D. F. M. (2009). The Brian simulator. Frontiers in Neuroscience 3, 192–197. doi:10.3389/neuro.01.026.2009

Gratiy, S. L., Billeh, Y. N., Dai, K., Mitelut, C., Feng, D., Gouwens, N. W., et al. (2018). Bionet: A python interface to neuron for modeling large-scale networks. PLoS One 13, e0201630

Guennebaud, G., Jacob, B., et al. (2010). Eigen v3. http://eigen.tuxfamily.org

Hagen, E., Næss, S., Ness, T. V., and Einevoll, G. T. (2018). Multimodal modeling of neural network activity: computing lfp, ecog, eeg, and meg signals with lfpy 2.0. Frontiers in neuroinformatics 12, 92

Hennessy, J. L. and Patterson, D. A. (2017). Computer Architecture, Sixth Edition: A Quantitative Approach (San Francisco, CA, USA: Morgan Kaufmann Publishers Inc.), 6th edn.

Hennessy, J. L. and Patterson, D. A. (2019). A new golden age for computer architecture. Communications of the ACM 62, 48–60. doi:10.1145/3282307

Heroux, M. A. (2015). Editorial: ACM TOMS Replicated Computational Results Initiative. ACM Transactions on Mathematical Software 41, 1–5. doi:10.1145/2743015

Hettrick, S., Antonioletti, M., Carr, L., Chue Hong, N., Crouch, S., De Roure, D., et al. (2014). Uk Research Software Survey 2014. doi:10.5281/ZENODO.14809

Hines, M. (1984). Efficient computation of branched nerve equations. International journal of bio-medical computing 15, 69–76

Hines, M., Davison, A., and Muller, E. (2009). NEURON and Python. Frontiers in Neuroinformatics 3, 1. doi:10.3389/neuro.11.001.2009

Hines, M., Kumar, S., and Schürmann, F. (2011a). Comparison of neuronal spike exchange methods on a blue gene/p supercomputer. Frontiers in computational neuroscience 5, 49

Hines, M., Kumar, S., and Schürmann, F. (2011b). Comparison of neuronal spike exchange methods on a Blue Gene/P supercomputer. Frontiers in computational neuroscience 5, 49. doi:10.3389/fncom.2011.00049

Hines, M. L. and Carnevale, N. T. (1997). The NEURON Simulation Environment. Neural Computation 9, 1179–1209. doi:10.1162/neco.1997.9.6.1179

Hines, M. L., Eichner, H., and Schürmann, F. (2008a). Neuron splitting in compute-bound parallel network simulations enables runtime scaling with twice as many processors. Journal of computational neuroscience 25, 203–210. doi:10.1007/s10827-007-0073-3

Hines, M. L., Markram, H., and Schürmann, F. (2008b). Fully implicit parallel simulation of single neurons. Journal of computational neuroscience 25, 439–448. doi:10.1007/s10827-008-0087-5

Hjorth, J. J. J., Kozlov, A., Carannante, I., Frost Nylén, J., Lindroos, R., Johansson, Y., et al. (2020). The microcircuits of striatum in silico. Proceedings of the National Academy of Sciences 117, 9554–9565. doi:10.1073/pnas.2000671117

HPE (2022). Hpe sgi 8600 system. https://support.hpe.com/hpesc/public/docDisplay?docId=emr_na-a00025339en_us. [Online; accessed 05-January-2022]

Jordan, J., Ippen, T., Helias, M., Kitayama, I., Sato, M., Igarashi, J., et al. (2018). Extremely scalable spiking neuronal network simulation code: From laptops to exascale computers. Frontiers in Neuroinformatics 12. doi:10.3389/fninf.2018.00002

Keating, S. M., Waltemath, D., König, M., Zhang, F., Dräger, A., Chaouiya, C., et al. (2020). Sbml level 3: an extensible format for the exchange and reuse of biological models. Molecular systems biology 16, e9110

Kumbhar, P., Awile, O., Keegan, L., Alonso, J. B., King, J., Hines, M., et al. (2020). An Optimizing Multi-platform Source-to-source Compiler Framework for the NEURON MODeling Language. In Computational Science – ICCS 2020, eds. V. V. Krzhizhanovskaya, G. Závodszky, M. H. Lees, J. J. Dongarra, P. M. A. Sloot, S. Brissos, and J. Teixeira (Cham: Springer International Publishing), Lecture Notes in Computer Science, 45–58. doi:10.1007/978-3-030-50371-0_4

Kumbhar, P., Hines, M., Fouriaux, J., Ovcharenko, A., King, J., Delalondre, F., et al. (2019). CoreNEURON: An Optimized Compute Engine for the NEURON Simulator. Frontiers in Neuroinformatics 13, 63. doi:10.3389/fninf.2019.00063

Kumbhar, P., Hines, M., Ovcharenko, A., Mallon, D. A., King, J., Sainz, F., et al. (2016). Leveraging a Cluster-Booster Architecture for Brain-Scale Simulations. In High Performance Computing, eds. J. M. Kunkel, P. Balaji, and J. Dongarra (Cham: Springer International Publishing), vol. 9697. 363–380. doi:10.1007/978-3-319-41321-1_19. Series Title: Lecture Notes in Computer Science

Lam, S. K., Pitrou, A., and Seibert, S. (2015). Numba: A llvm-based python jit compiler. In Proceedings of the Second Workshop on the LLVM Compiler Infrastructure in HPC (New York, NY, USA: Association for Computing Machinery), LLVM ‘15. doi:10.1145/2833157.2833162

Lee, V. W., Kim, C., Chhugani, J., Deisher, M., Kim, D., Nguyen, A. D., et al. (2010). Debunking the 100X GPU vs. CPU myth: an evaluation of throughput computing on CPU and GPU. ACM SIGARCH Computer Architecture News 38, 451–460. doi:10.1145/1816038.1816021

Leloup, J.-C., Gonze, D., and Goldbeter, A. (1999). Limit cycle models for circadian rhythms based on transcriptional regulation in drosophila and neurospora. Journal of biological rhythms 14, 433–448

Lindén, H., Hagen, E., Leski, S., Norheim, E. S., Pettersen, K. H., and Einevoll, G. T. (2014). Lfpy: a tool for biophysical simulation of extracellular potentials generated by detailed model neurons. Frontiers in neuroinformatics 7, 41

Lytton, W. W., Seidenstein, A. H., Dura-Bernal, S., McDougal, R. A., Schürmann, F., and Hines, M. L. (2016). Simulation Neurotechnologies for Advancing Brain Research: Parallelizing Large Networks in NEURON. Neural Computation 28, 2063–2090. doi:10.1162/NECO_a_00876

Malik, R., Dougherty, K. A., Parikh, K., Byrne, C., and Johnston, D. (2016). Mapping the electrophysiological and morphological properties of ca 1 pyramidal neurons along the longitudinal hippocampal axis. Hippocampus 26, 341–361

Markram, H., Muller, E., Ramaswamy, S., Reimann, M., Abdellah, M., Sanchez, C., et al. (2015). Reconstruction and Simulation of Neocortical Microcircuitry. Cell 163, 456–492. doi:10.1016/j.cell.2015.09.029

McDougal, R. A., Bulanova, A. S., and Lytton, W. W. (2016). Reproducibility in computational neuroscience models and simulations. IEEE Transactions on Biomedical Engineering 63, 2021–2035

McDougal, R. A., Conte, C., Eggleston, L., Newton, A. J. H., and Galijasevic, H. (2022). Efficient Simulation of 3D Reaction-Diffusion in Models of Neurons and Networks. Frontiers in Neuroinformatics 16

McDougal, R. A., Hines, M. L., and Lytton, W. W. (2013). Reaction-diffusion in the neuron simulator. Frontiers in neuroinformatics 7, 28

McDougal, R. A., Morse, T. M., Carnevale, T., Marenco, L., Wang, R., Migliore, M., et al. (2017). Twenty years of ModelDB and beyond: Building essential modeling tools for the future of neuroscience. Journal of Computational Neuroscience 42, 1–10. doi:10.1007/s10827-016-0623-7

Metzner, C., Mäki-Marttunen, T., Karni, G., McMahon-Cole, H., and Steuber, V. (2020). The effect of alterations of Schizophrenia-Associated genes on gamma band oscillations

Meurer, A., Smith, C. P., Paprocki, M., Certík, O., Kirpichev, S. B., Rocklin, M., et al. (2017). SymPy: Symbolic computing in Python. PeerJ Computer Science 3, e103. doi:10.7717/peerj-cs.103

Meyer, M. (2014). Continuous Integration and Its Tools. IEEE Software 31, 14–16. doi:10.1109/MS.2014.58

Migliore, M., Cannia, C., Lytton, W. W., Markram, H., and Hines, M. L. (2006). Parallel network simulations with NEURON. Journal of Computational Neuroscience 21, 119–129. doi:10.1007/s10827-006-7949-5

Migliore, M., Cavarretta, F., Hines, M. L., and Shepherd, G. M. (2014). Distributed organization of a brain microcircuit analyzed by three-dimensional modeling: the olfactory bulb. Frontiers in Computational Neuroscience 8, 50. doi:10.3389/fncom.2014.00050

Miller, G. (2006). A Scientist’s Nightmare: Software Problem Leads to Five Retractions. Science 314, 1856–1857. doi:10.1126/science.314.5807.1856

Muller, E., Bednar, J. A., Diesmann, M., Gewaltig, M.-O., Hines, M., and Davison, A. P. (2015). Python in neuroscience. Frontiers in Neuroinformatics 9

Mulugeta, L., Drach, A., Erdemir, A., Hunt, C. A., Horner, M., Ku, J. P., et al. (2018). Credibility, replicability, and reproducibility in simulation for biomedicine and clinical applications in neuroscience. Front. Neuroinform. 12, 18

Neely, J., de Supinski, B. R., and Still, C. H. (2017). Application modernization for the exascale era. Computing in Science & Engineering 19, 6–8. doi:10.1109/MCSE.2017.3421548

Newton, A. J., McDougal, R. A., Hines, M. L., and Lytton, W. W. (2018). Using neuron for reaction-diffusion modeling of extracellular dynamics. Frontiers in neuroinformatics 12, 41

Newton, T. H., Reimann, M. W., Abdellah, M., Chevtchenko, G., Muller, E. B., and Markram, H. (2021). In silico voltage-sensitive dye imaging reveals the emergent dynamics of cortical populations. Nature Communications 12, 3630. doi:10.1038/s41467-021-23901-7

Neymotin, S. A., Daniels, D. S., Caldwell, B., McDougal, R. A., Carnevale, N. T., Jas, M., et al. (2020). Human neocortical neurosolver (hnn), a new software tool for interpreting the cellular and network origin of human meg/eeg data. Elife 9, e51214

Pham, D.-T. J., Yu, G. J., Bouteiller, J.-M. C., and Berger, T. W. (2021). Bridging hierarchies in multi-scale models of neural systems: Look-up tables enable computationally efficient simulations of non-linear synaptic dynamics. Frontiers in computational neuroscience, 88

Pimentel, J. M., Moioli, R. C., de Araujo, M. F. P., Ranieri, C. M., Romero, R. A. F., Broz, F., et al. (2021). Neuro4PD: An initial neurorobotics model of parkinson’s disease. Front. Neurorobot. 15, 640449

Pronold, J., Jordan, J., Wylie, B. J. N., Kitayama, I., Diesmann, M., and Kunkel, S. (2022). Routing Brain Traffic Through the Von Neumann Bottleneck: Parallel Sorting and Refactoring. Frontiers in Neuroinformatics 15

Ranieri, C. M., Pimentel, J. M., Romano, M. R., Elias, L. A., Romero, R. A. F., Lones, M. A., et al. (2021). A Data-Driven biophysical computational model of parkinson’s disease based on marmoset monkeys. IEEE Access 9, 122548–122567

Reimann, M. W., Nolte, M., Scolamiero, M., Turner, K., Perin, R., Chindemi, G., et al. (2017). Cliques of Neurons Bound into Cavities Provide a Missing Link between Structure and Function. Frontiers in Computational Neuroscience 11, 48. doi:10.3389/fncom.2017.00048

Reuther, A., Michaleas, P., Jones, M., Gadepally, V., Samsi, S., and Kepner, J. (2019). Survey and Benchmarking of Machine Learning Accelerators. In 2019 IEEE High Performance Extreme Computing Conference (HPEC) (Waltham, MA, USA: IEEE), 1–9. doi:10.1109/HPEC.2019.8916327

Romaro, C., Najman, F. A., Lytton, W. W., Roque, A. C., and Dura-Bernal, S. (2021). NetPyNE implementation and scaling of the Potjans-Diesmann cortical microcircuit model. Neural Comput. 33, 1993–2032

Salmon, J. K., Moraes, M. A., Dror, R. O., and Shaw, D. E. (2011). Parallel random numbers: As easy as 1, 2, 3. In Proceedings of 2011 International Conference for High Performance Computing, Networking, Storage and Analysis (New York, NY, USA: Association for Computing Machinery), SC ‘11, 1–12. doi:10.1145/2063384.2063405

Schirner, M., Domide, L., Perdikis, D., Triebkorn, P., Stefanovski, L., Pai, R., et al. (2022). Brain simulation as a cloud service: The virtual brain on ebrains. NeuroImage, 118973

Sekiguchi, K., Medlock, L., Dura-Bernal, S., Prescott, S. A., and Lytton, W. W. (2021). Multiscale computer model of the spinal dorsal horn reveals changes in network processing associated with chronic pain

Sivagnanam, S., Gorman, W., Doherty, D., Neymotin, S. A., Fang, S., Hovhannisyan, H., et al. (2020). Simulating large-scale models of brain neuronal circuits using google cloud platform. In Practice and Experience in Advanced Research Computing (New York, NY, USA: Association for Computing Machinery), PEARC ‘20, 505–509

Sivagnanam, S., Majumdar, A., Yoshimoto, K., Astakhov, V., Bandrowski, A. E., Martone, M. E., et al. (2013). Introducing the neuroscience gateway. IWSG 993

Stimberg, M., Brette, R., and Goodman, D. F. (2019). Brian 2, an intuitive and efficient neural simulator. eLife doi:10.7554/eLife.47314

Tikidji-Hamburyan, R. A., Narayana, V., Bozkus, Z., and El-Ghazawi, T. A. (2017). Software for Brain Network Simulations: A Comparative Study. Frontiers in Neuroinformatics 11, 46. doi:10.3389/fninf.2017.00046

Van Geit, W., Gevaert, M., Chindemi, G., Rössert, C., Courcol, J.-D., Muller, E. B., et al. (2016). BluePyOpt: Leveraging Open Source Software and Cloud Infrastructure to Optimise Model Parameters in Neuroscience. Frontiers in Neuroinformatics 10. doi:10.3389/fninf.2016.00017

Virtanen, P., Gommers, R., Oliphant, T. E., Haberland, M., Reddy, T., Cournapeau, D., et al. (2020). Scipy 1.0: fundamental algorithms for scientific computing in python. Nature methods 17, 261–272

Volk, V. L., Hamilton, L. D., Hume, D. R., Shelburne, K. B., and Fitzpatrick, C. K. (2021). Integration of neural architecture within a finite element framework for improved neuromusculoskeletal modeling. Sci. Rep. 11, 22983

Willenbring, J. M. (2015). Replicated Computational Results (RCR) Report for “BLIS: A Framework for Rapidly Instantiating BLAS Functionality”. ACM Transactions on Mathematical Software 41, 1–4. doi:10.1145/2738033

Wolfe, M. (2021). Performant, Portable, and Productive Parallel Programming With Standard Languages. Computing in Science & Engineering 23, 39–45. doi:10.1109/MCSE.2021.3097167

